# Ovarian Cancer Risk Variants are Enriched in Histotype-Specific Enhancers that Disrupt Transcription Factor Binding Sites

**DOI:** 10.1101/2020.02.21.960468

**Authors:** Michelle R. Jones, Pei-Chen Peng, Simon G. Coetzee, Jonathan Tyrer, Alberto L. Reyes, Rosario I. Corona de la Fuente, Brian Davis, Stephanie Chen, Felipe Dezem, Ji-Heui Seo, Ovarian Cancer Association Consortium, Benjamin P. Berman, Matthew L. Freedman, Jasmine T. Plummer, Kate Lawrenson, Paul Pharoah, Dennis J. Hazelett, Simon A. Gayther

## Abstract

Quantifying the functional effects of complex disease risk variants can provide insights into mechanisms underlying disease biology. Genome wide association studies (GWAS) have identified 39 regions associated with risk of epithelial ovarian cancer (EOC). The vast majority of these variants lie in the non-coding genome, suggesting they mediate their function through the regulation of gene expression by their interaction with tissue specific regulatory elements (REs). In this study, by intersecting germline genetic risk data with regulatory landscapes of active chromatin in ovarian cancers and their precursor cell types, we first estimated the heritability explained by known common low penetrance risk alleles. The narrow sense heritability 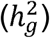 of both EOC overall and high grade serous ovarian cancer (HGSOCs) was estimated to be 5-6%. Partitioned SNP-heritability across broad functional categories indicated a significant contribution of regulatory elements to EOC heritability. We collated epigenomic profiling data for 77 cell and tissue types from public resources (Roadmap Epigenomics and ENCODE), and H3K27Ac ChIP-Seq data generated in 26 ovarian cancer-relevant cell types. We identified significant enrichment of risk SNPs in active REs marked by H3K27Ac in HGSOCs. To further investigate how risk SNPs in active REs influence predisposition to ovarian cancer, we used motifbreakR to predict the disruption of transcription factor binding sites. We identified 469 candidate causal risk variants in H3K27Ac peaks that break TF motifs (enrichment P-Value < 1×10^−5^ compared to control variants). The most frequently broken motif was REST (P-Value = 0.0028), which has been reported as both a tumor suppressor and an oncogene. These systematic functional annotations with epigenomic data highlight the specificity of the regulatory landscape and demonstrate functional annotation of germline risk variants is most informative when performed in highly relevant cell types.

## Introduction

Epithelial ovarian cancer (EOC) consists of five histological subtypes of invasive disease; High grade serous (HGSOC), low grade serous (LGSOC), mucinous (MOC), endometrioid (EnOC) and clear cell (CCOC) ovarian cancer. The majority of serous cases are diagnosed at a late stage and this contributes to the poor prognosis and resistance to standard chemotherapeutic treatments frequently observed^1–3^. Ovarian tumors of low malignant potential (LMP) comprise ∼20% of cases and only a small minority will progress to invasive disease. Each histotype shows differences in underlying biology, genetic risk and to some extent different epidemiological and lifestyle risk factors. They may also derive from different cell types, with fallopian tube secretory epithelial cells the likely cell of origin for most serous tumors ^4,5^, and endometriosis the putative precursor of CCOC and EnOC ^6–8^. Uncovering the underlying genetic architecture of different EOC histotypes is an urgent need and may be the most effective approach to reduce mortality due to EOC ^9^.

Less than forty percent of the estimated narrow sense heritability of ovarian cancer is explained by known coding pathogenic mutations in susceptibility genes including *BRCA1, BRCA2, BRIP1, RAD51C* and *RAD51D* ^10^. Genome wide association studies (GWAS) have identified 40 independent regions associated with EOC risk ^11^. Some regions are associated with specific histotypes, while others appear pleiotropic across different EOC histotypes ^11–21^ or other phenotypes (e.g. breast cancer) ^21,22^. Combined, these common, low risk alleles explain a fraction of the narrow sense heritability for ovarian cancer. Heritability estimates are complicated by linkage disequilibrium, which often results in the identification of tens to hundreds of tightly correlated SNPs at each susceptibility locus ^23^.

The vast majority of risk alleles for common complex traits identified by GWAS lie in the non-protein coding DNA regions with their mechanisms of function largely unknown ^24^. Several studies of complex disease phenotypes have shown that risk variants are enriched in regulatory elements, suggesting that they function through the differential regulation of gene expression ^25–28^. Many regulatory elements can be identified by epigenomic modifications; for example, H3K4me1 and H3K4me2 histone modifications correlate with poised enhancers, H3K27Ac with active enhancers and CTCF with gene repressors or the flanking boundaries of topologically associated domains ^29–32^. Publicly available resources such as the Encyclopedia of DNA Elements (ENCODE) and the Roadmap Epigenome Mapping Consortium (REMC) have characterized the epigenomic architecture of a multitude of cell types, showing that the epigenomes and transcriptional program are highly tissue-specific ^29,33^. Analyses of acetylated lysine 27 of histone H3 (H3K27Ac) in primary tissues shows that >80% of cell type-specific regulatory elements lie in putative enhancers, reinforcing previous observations that cell type-specific enhancers drive the spatial and temporal diversity of gene expression ^29,34^.

We hypothesize that common ovarian cancer risk SNPs are located within tissue specific regulatory elements and are likely to function by altering the activity of enhancers active in ovarian cancers and cell types that represent likely precursors of the different EOC histotypes. We applied systematic computational approaches to identify regulatory elements that are potentially disrupted at EOC GWAS risk loci. We first estimated the heritability for each EOC histotype using common SNPs, taking into account linkage disequilibrium; and then partitioned narrow sense heritability across general broad functional categories. We focused our analyses on 40 germline GWAS risk loci previously reported for one or more EOC histotypes with the aim of identifying putative regulatory elements and transcription factors associated with EOC risk variants and the initiation and development of EOC.

## Methods

### Genotyping datasets for ovarian cancer

Summary statistics were available from the largest published meta-analysis of 25,509 EOC cases and 40,941 controls ^11^. This analysis included EOC cases from the five major histotypes of invasive disease; HGSOC; n = 13,037, LGSOC; n = 1,012, MOC; n = 1,149, EnOC; n = 2,764 and CCOC; n = 1,366, and borderline serous; n = 1,954 and EOC cases of either unknown or undefined histology (n = 2,749). This analysis utilized genotypes based on the 1000 Genomes Project reference panel of 11,403,952 common variants (MAF>1%).

We further curated all previously reported genome-wide significant risk regions for EOC (including all histotypes) to identify a credible causal set of SNPs at each locus for all invasive ovarian cancer and for each histotype where there was evidence of a risk association ^9–19^. This identified 39 risk regions for different histotypes at genome wide significance (P<5.0 x 10^−8^) (Supplementary Table 1). Variant position and rsid for each variant were validated in dbSNP146 with hg19/GRCh37 coordinates.

### Epigenomic and datasets for ovarian cancer and their precursor tissues

Publicly available epigenomic profiling datasets were collected from the Roadmap Epigenomics Mapping Consortium ^34^ and the ENCODE project ^29^ (labelled as ‘ENCODE2012’ in this study, Supplementary Table 2). Additionally, a collection of chromatin immunoprecipitation-sequencing (ChIP-seq) for H3K27Ac in ovarian cancer related cell and tissue types that were generated in house was compiled. This includes precursor normal and ovarian cancer cell lines from previously published studies and newly generated H3K27Ac ChIP-Seq in additional cell lines and primary tumors (Supplementary Table 3). Briefly, we have generated H3K27Ac-ChIP-seq data for: Twenty primary EOC tumors, five each for the different histotypes of invasive ovarian cancer (HGSOC, CCOC, EnOC and MOC) (Supplementary Table 3); twelve established EOC cell lines that model; undifferentiated EOC (HEYA8), HGSOC (CaOV3, UWB1.289, Kuramochi, OVCA429), LGSOC (VOA1056, OAW42), CCOC (JHOC5, ES2 and RMG-II) and MOC (GFTR230, MCAS, EFO27); and three ovarian cancer precursor cell types; fallopian tube secretory epithelial cells ((FTSECs), FT246, FT33), ovarian surface epithelial cells ((OSECs), IOSE4 and IOSE11) and endometrioid epithelial cells (EEC16) ^35^ (Supplementary Table 3). Methods for H3K27Ac-ChIP-seq and peak calling that was previously published have been described ^11,29,36–41^.

H3K27Ac ChIP-Seq for six new cell lines (EFO27, VOA1056, HEYA8, Kuramochi, ES2, RMG-II) was performed according to previously published methods ^42^. Peak calling was performed using the AQUAS pipeline ^43^. Reads were aligned against the reference human genome hg38. Quality control metrics were computed for each individual replicate, including number of reads, percentage of duplicate reads, normalized strand coefficient, relative strand correlation and fraction of reads in called peaks. Two biological replicates were available for EFO27, VOA1056 and Kuramochi. Peak calling was performed with macs2 with pooled replicate peaks that overlap 50% or more in each individual replicate selected for the final peak set. When replicates were not available (HEYA8, ES2 and RMG-II) psuedo replicates were formed and pooled peaks selected in the same manner from these pseudo replicates. To create consensus peak sets across a single histotype for enrichment analyses, peaks with least 50% overlap with at least one other peak in two or more samples from a histotype group were retained, with the boundaries stretched to the edge of each peak in the overlap. Files were then concatenated and peak co-ordinates merged such that if records within the concatenated file were overlapping they were combined into a single peak.

We generated chromatin state calls in REMC and ENCODE2012 samples using StatepaintR ^44^ (Supplementary Table 2). This approach uses human expert rule-based segmentations, which allows the user to designate combinations of epigenomic marks to represent functional chromatin states. StatepaintR annotates chromatin states based upon available epigenomic marks, accommodating for the practical situation that not all histone marks are available for all samples. These chromatin state annotations are also released in the StateHub Model Repository under TrackHub ID 5813b67f46e0fb06b493ceb0 (www.statehub.org/).

### Estimation of SNP-heritability

We estimated the variance explained by known SNP effects, or SNP-heritability, by using linkage disequilibrium score regression (LDSC) ^45,46^, version 1.0.0. LDSC models the expected *χ*^2^ statistics from a GWAS of SNP *j* as

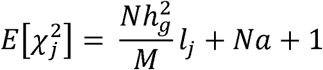

where *N* is the number of individuals; *M* is the number of SNPs, such that 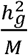 is the average heritability explained per SNP; *α* is a constant measuring the contribution of confounding biases, such as cryptic relatedness and population stratification; *l*_*j*_ is the LD score of SNP *j* defined as 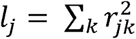, where 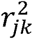 is the Pearson correlation between SNP *j* and SNP *k*, and *k* denotes other SNPs within the LD region. The LD scores were pre-calculated from phased European-ancestry individuals from the 1000 Genomes Project reference panel v3 ^45^.

### Partitioning SNP-heritability into functional categories

To examine the importance of specific functional categories in SNP-heritability, we applied stratified LD score regression ^46^ to EOC and HGSOC GWAS summary statistics. The goal was to partition SNP-heritability into functional categories by combining SNPs in the same LD region together and quantify their overlaps with regions of interest. The stratified LDSC model was adapted from the above-mentioned regular LDSC model:

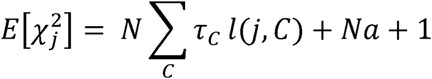

where *C* represents the functional categories; *τ*_*C*_ denotes the per-SNP contribution to heritability of category *C*; *l* (*j, C*) is the LD score of SNP *j* falling in category *C*, calculated as 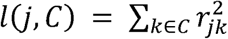; all the other parameters are the same as in LDSC. The category-specific enrichment was defined as the proportion of SNP-heritability in the category divided by the proportion of SNPs in the same category.

The partitioned-heritability analyses were performed with two different sets of functional categories. The first is a full baseline model with 24 general broad functional annotations from public datasets, which were inclusive of all publicly available cell types and post-processed in Gusev et al. ^47^. The 24 annotations include coding, 3’UTR, 5’UTR, promoter, and intron regions from UCSC Genome Browser ^47,48^; regions conserved in mammals ^49,50^; combined chromHMM and Segway predictions comprising CTCF-bound regions, promoter-flanking, transcribed, transcription start site (TSS), strong enhancer, weak enhancer, repressed annotations ^51^; digital genomic footprint (DGF) and transcription factor binding sites (TFBS) from ENCODE ^47^; open chromatin regions as reflected by DNase I hypersensitivity sites (DHSs) from a union of all cell types and a union of only fetal cell types on ENCODE and Roadmap Epigenomics ^52^; FANTOM5 enhancer ^53^; H3K27Ac ^54,55^, H3K4me1 ^29^, and H3K4me3 ^29^ histone marks from a union over cell types on Roadmap Epigenomics; super-enhancers obtained from ^55^.

The second set contains 15 cell-type-specific annotations for H3K27Ac marks, which represent precursor normal and ovarian cancer cell lines (see the ‘Epigenomic profiling’ section for details). We added these cell-type-specific annotations individually to the full baseline model, which resulted in 15 models for EOC and 15 models for HGSOC. This cell-type-specific analysis helped measure how much more the annotation contributes on top of the rest of the full baseline model, and to justify which cell type is more enriched than the others.

### Enrichment of credible causal SNPs in biofeatures

EOC credible causal risk variants were combined to create the full credible set (n=1432), and then split to represent sets of risk variants associated with each EOC histotypes. The background set of variants used in functional annotation and enrichment analysis were generated by aggregating SNPs within 2Mb (1Mb +/-) of the credible causal set, in an attempt to maintain similar genetic architecture (e.g. linkage disequilibrium) as credible causal risk variants. Functional annotation of credible causal SNPs was performed with SNPnexus ^56^ using SIFT ^57^ and Polyphen ^58^ for protein effect, ENCODE ^29^, Roadmap Epigenomics ^34^, and Ensembl Regulatory Build ^59^ for regulatory elements, and CADD ^60^, DeepSEA ^61^ and FunSeq2 ^62^ for non-coding variation scoring. The difference between the average FunSeq2 functional score for the foreground and background SNP lists was determined with a two tailed t test.

Enrichment analysis was performed with the FunciVar package (https://github.com/Simon-Coetzee/funcivar), a tool for annotation and functional enrichment of variant sets. In principle, FunciVar first takes two lists of variants as inputs: 1) a list of target variants, in this analysis the credible causal set of risk SNPs, which act as the foreground, and 2) a list of control variants, which act as the background. The background SNP lists from each locus were combined as necessary to ensure the local background set of variants for each locus was included in the histotype-specific enrichment. FunciVar then intersects each variant with biofeatures, which were provided as bed files. The likelihood of true enrichment for each variant list is modeled under the beta-binomial distribution.

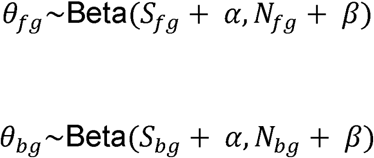

where *S* is the number of observed overlaps with biofeature, *N* is the number of total variants, and subscripts *bg* and *fg* denote background and foreground respectively. FunciVar uses an uninformative Jeffreys prior, which set *α* = 0.5 and *β* = 0.5. To estimate the true enrichment, FunciVar by default simulates 10,000 times to obtain a distribution of foreground enrichment probability, *θ* _*fg*_, and a distribution of background enrichment probability, *θ* _*bg*_. The two sets of simulated probabilities were next directly subtracted to obtain a distribution of differences. FunciVar calculates a 95% credible interval for the range of enrichment probability differences between the two lists of variants. Enrichment is reported as the median of this credible interval, within the range of −1 to 1, where 1 means strong enrichment and −1 means strong depletion. The significance of results is reported as probability that foreground SNPs have more overlaps with the biofeature than background SNPs, within the range of 0 to 1, the higher the more confident. Results are plotted with significantly enriched biofeatures shown in color, and non-significantly enriched biofeatures shown in grey.

### Identifying transcription factor binding consequences of EOC credible causal variants in enhancers

To identify the potential consequences of EOC risk variants in EOC enhancers we used MotifBreakR ^63^ to predict the transcription factor binding sites that a variant disrupts and the extent of disruptiveness. MotifBreakR uses a position weight matrix to score the difference of binding between reference and alternative alleles for every possible window that includes the variant, and then categorizes the normalized difference score as effect of the target variant (strong, weak, or neutral). We used seven TFBS motif databases; ENCODE motifs ^64^, Factorbook ^65^, Hocomoco ^66^, Homer ^67^, Transfac ^68^, Jaspar ^69^ and MotifDb ^70^.

To identify significant TFs that were predicted to be impacted by the alternate allele at credible causal variants, we applied FunciVar package again. We curated two lists: 1) the foreground list, which are credible causal variants that intersect H3K27Ac peaks in any EOC cell type, and 2) the background list; credible causal variants that did not intersect H3K27Ac peaks in any EOC cell type. Significant differences in likelihood of the alternate allele of a credible causal variant disrupting a TFBS are reported for each TF.

## Results

### Regulatory elements significantly account for ovarian cancer heritability

The aim was to evaluate the functional significance of common, genetic variants associated with epithelial ovarian cancer (EOC) risk identified by GWAS, and the contribution of different functional states to EOC heritability. We utilized genotype data pooled from multiple GWAS comprising 25,509 EOC cases and 40,941 controls stratified into five major histotypes of invasive or low grade/borderline disease: High grade serous (HGSOC), low grade serous (LGSOC), mucinous (MOC) endometrioid (EnOC) and clear cell (CCOC) ovarian cancers (see Methods). These analyses identified thirty-nine different risk regions at P<5 x 10^−8^ either for all invasive EOC or specific to different histotypes. Fine mapping of these regions identified a total of 1,432 credible risk variants at these loci, ranging from 3 to 192 risk SNPs per region (Supplementary Table 1).

We estimated the variance explained by known SNP effects, or SNP-heritability, using linkage disequilibrium score regression (LDSC) ^45,46^. LDSC measures narrow sense heritability (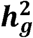, ‘SNP-heritability’ henceforth) using GWAS summary statistics to explicitly model linkage disequilibrium. Estimates of SNP-heritability ranged from nearly 0 - 6% for the different EOC histotypes (Figure 1), with the highest heritability explained by risk variants associated with the HGSOC histotype and the lowest heritability for risk variants associated with LGSOC.

**Figure 1.**
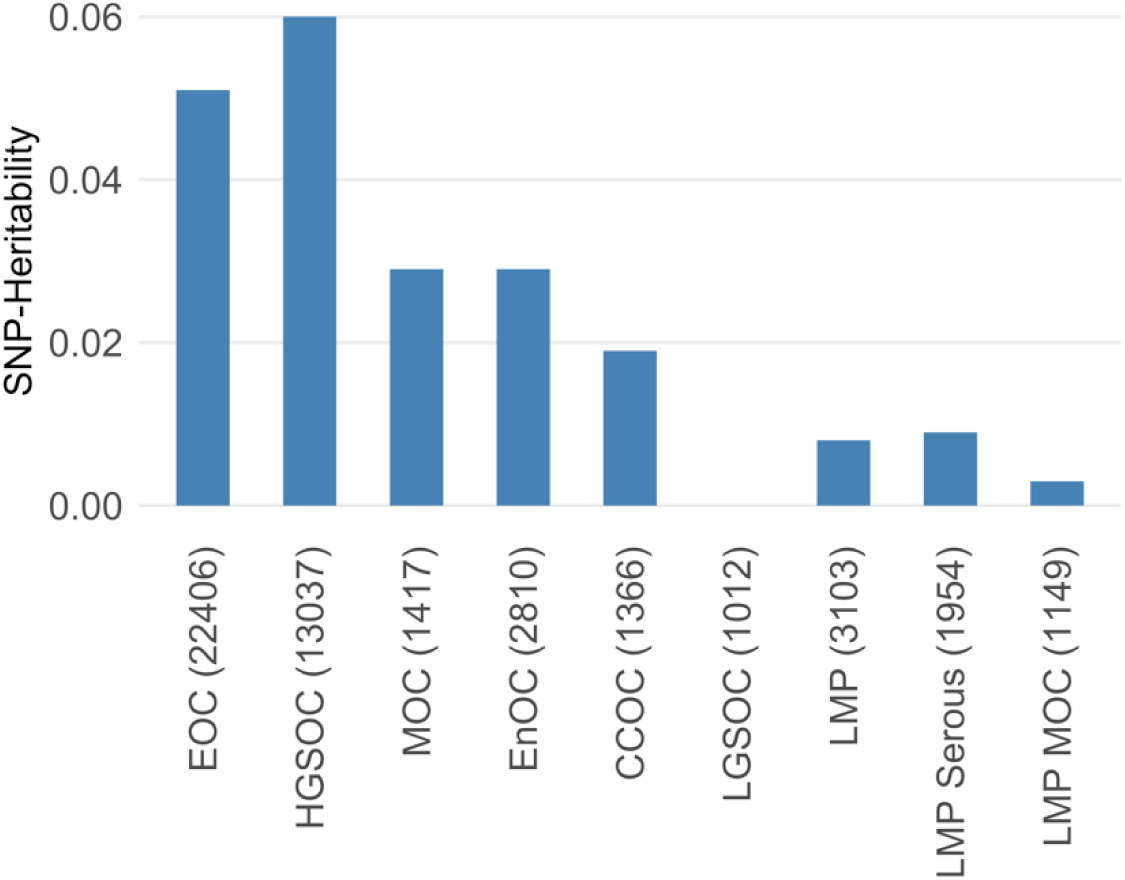
Estimates of SNP-heritability 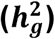 explained by common SNPs. Overall SNP heritability calculated based on GWAS summary statistics for each EOC histotype. The GWAS included 40,941 control cases and the number of cases by histotypes are shown in parentheses. EOC: Epithelial ovarian cancer; HGSOC: high grade serous ovarian cancer; MOC: mucinous ovarian cancer; EnOC: endometrioid ovarian cancer; CCOC: clear cell ovarian cancer; LGSOC: low grade serous ovarian cancer; LMP: low malignant potential

Next, we partitioned SNP-heritability across 24 broad non-cell-type-specific ‘functional’ categories (see Methods) ^71^. For these analyses, EOC cases were stratified into two group - ‘all invasive EOC’ and HGSOC - based on the results of heritability analyses (Figure 1). We observed a significant contribution of several functional features that may regulate gene expression to EOC heritability (Table 1). For example, 27% of 1,432 candidate causal risk variants coincided with the histone modification H3K27Ac, accounting for 97% of the estimated SNP-heritability (3.6-fold enrichment, P-Value = 0.006). Other significant functional elements included 3 prime untranslated regions (3’UTR) (17.3-fold enrichment, P-Value = 0.015); promoters (8.7-fold enrichment, P-Value = 0.016); and super-enhancers (2.1-fold enrichment, P-Value = 0.02) (Table 1). HGSOC heritability was most strongly driven by 3’UTRs (18.4-fold enrichment, P-Value = 0.009) and H3K27Ac marks (1.8-fold enrichment, P-Value = 0.033).

**Table 1.** Enrichment estimates for 24 non-cell-type-specific functional categories for EOC and HGSOC. Enrichment was calculated as 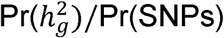, which shows the proportion of estimated SNP-heritability explained by the proportion of SNPs in the functional category. Statistically significant associations (P-values < 0.05) are marked in bold.

| Functional Category | All EOC |  | HGSOC |  |
| --- | --- | --- | --- | --- |
|  | Enrichment<br>t | P-value | Enrichment<br>t | P-value |
| 3'UTR | **17.29** | **0.02** | **18.40** | **0.01** |
| 5'UTR | -0.62 | 0.89 | 5.12 | 0.71 |
| Coding | 3.55 | 0.75 | 8.05 | 0.30 |
| Conserved | 24.94 | 0.06 | 21.82 | 0.07 |
| CTCF | -9.62 | 0.11 | -3.86 | 0.47 |
| DGF | 1.71 | 0.82 | -0.92 | 0.52 |
| DHS | 3.42 | 0.35 | 0.50 | 0.83 |
| Enhancer | 2.66 | 0.69 | 2.56 | 0.71 |
| FANTOM5 enhancer | -2.46 | 0.86 | 12.42 | 0.51 |
| Fetal DHS | 0.41 | 0.87 | -1.52 | 0.50 |
| H3K27Ac (Hnisz et al.) | **1.96** | **0.01** | **1.77** | **0.03** |
| H3K27Ac (PGC2) | **3.61** | **0.01** | 2.42 | 0.16 |
| H3Kme1 | 2.02 | 0.16 | 1.82 | 0.28 |
| H3Kme3 | 3.91 | 0.07 | 1.01 | 0.99 |
| H3K9ac | 3.25 | 0.33 | 0.90 | 0.96 |
| Intron | 1.46 | 0.12 | 1.24 | 0.35 |
| Promoter | **8.69** | **0.02** | 6.99 | 0.06 |
| Promoter flanking | 10.38 | 0.49 | -2.90 | 0.76 |
| Repressed | -0.32 | 0.07 | 0.67 | 0.64 |
| Superenhancer | **2.09** | **0.02** | 1.83 | 0.12 |
| Transcription factor binding site | 4.93 | 0.16 | 2.21 | 0.66 |
| Transcribed | 1.87 | 0.24 | 1.05 | 0.95 |
| TSS | 5.28 | 0.55 | 0.56 | 0.96 |
| Weak enhancer | 9.06 | 0.35 | 4.69 | 0.68 |

### Enrichment of EOC risk variants with different chromatin states by cell type

We integrated 1,432 credible causal risk variants with epigenomic data to evaluate enrichment of EOC risk variants in different chromatin states by cell type. We first focused on publicly available data from Roadmap Epigenomics and ENCODE which are mainly for non-ovarian epigenomic datasets. We annotated the full credible set of EOC risk SNPs with SNPnexus ^72^ to map each variant to intergenic, intronic, 3’ or 5’ UTR or exonic regions (Supplementary Figure 1a). The majority of credible causal SNPs (96%) fall into non-protein coding DNA regions; 71% of SNPs lie in intergenic regions; and 25% of SNPs lie in intronic regions. We obtained a functional impact score for each variant through FunSeq2 scoring algorithms ^62^. The average functional impact score of EOC risk variants was 0.404, which is significantly higher than regional, matched background SNPs (0.2404; P-Value = 2.02×10^−49;^ Supplementary Figure 1b).

We performed enrichment analyses to test whether EOC risk SNPs are enriched within specific classes of biofeatures. We used StatePaintR ^44^ to combine epigenomic marks into chromatin state calls that represent functional elements, including active, poised, silenced, and weak states of enhancers and promoters. We first evaluated enrichment of EOC risk SNPs with chromatin states from Roadmap Epigenomics and ENCODE for publicly available tissues ^29,34^. Enrichment tests were performed using FunciVar (see Methods). Overall, we observed the greatest enrichment of EOC risk SNPs in active regulatory regions in digestive, immune, epithelial, liver, thymus, smooth muscle and stem cell types and each of the cancer-associated ENCODE2012 cell lines, which are all closely related cell types (Figure 2, Supplementary Table 4). In contrast, we observed a depletion of EOC risk SNPs in heterochromatin in 68 cell types, and an enrichment in polycomb repressed silenced regions in 48 cell types. Overall these analyses indicate that the enrichment of EOC risk SNPs in active regulatory regions is typically more cell-type restricted than in silenced regions.

**Figure 2.**
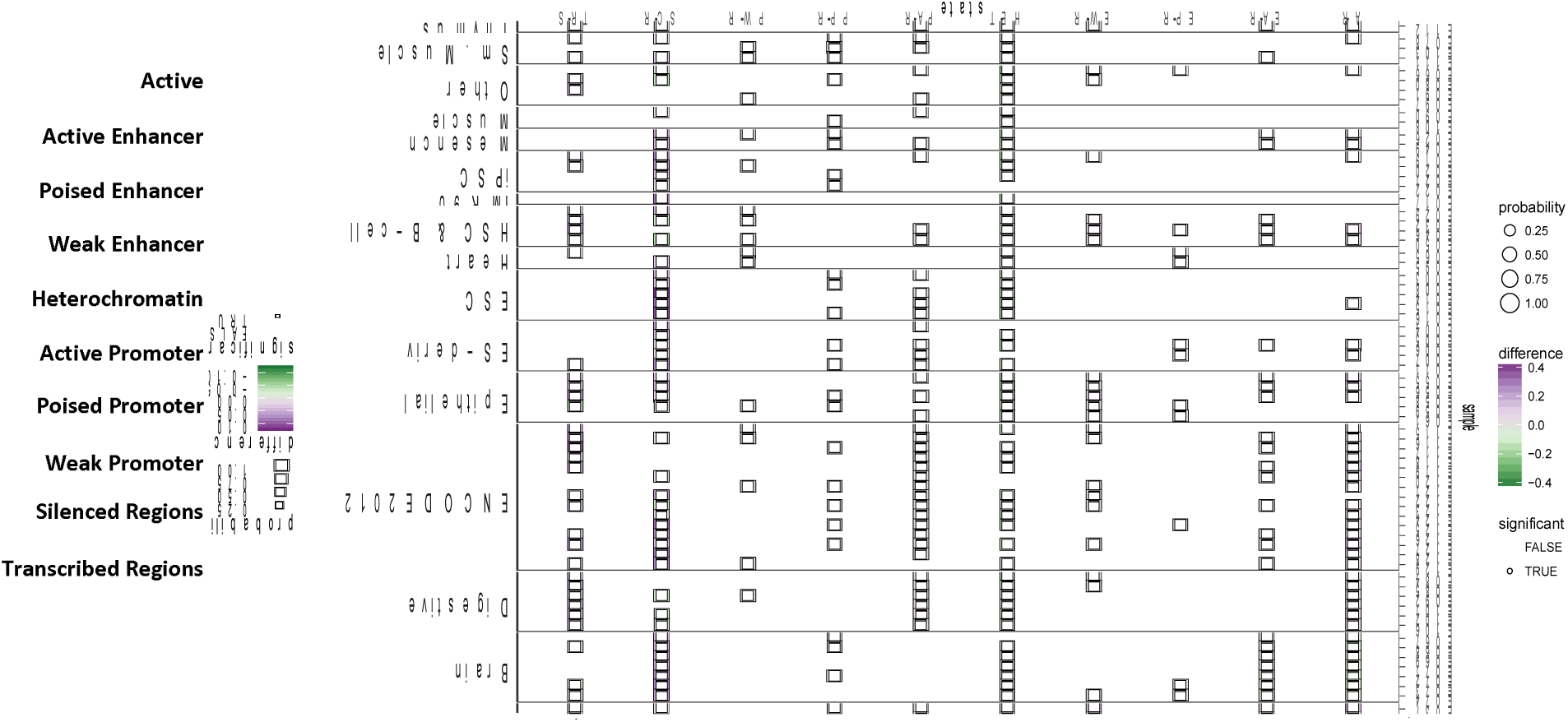
EOC risk variants are enriched in active regulatory elements. Enrichment analyses were performed in different chromatin states in REMC and ENCODE tissues and cell lines. Enriched biofeatures are shown in purple, depleted biofeatures in green, and non-significantly enriched biofeatures in grey. The size of the circle indicates the degree of confidence. EOC risk SNPs are significantly enriched in active regulatory elements in blood and T cells, digestive cell types and ENCODE cell lines.

We observed the strongest enrichment in an active regulatory chromatin state in stimulated primary T helper cells (E041) and primary T helper memory cells (E037), where 165 and 128 of 1432 EOC risk SNPs respectively overlapped active regions (Figure 2, Supplementary Table 5). There was also enrichment in active regulatory regions in all digestive tissue types (sigmoid colon, rectal mucosa, small intestine and stomach). By contrast, we found no evidence of enrichment for EOC risk SNPs in active regulatory regions in brain, heart or lung tissues, but instead observed enrichment for silenced regions in these tissue types.

### Enrichment of EOC risk variants in regions marked by H3K27Ac peaks in ovarian and non-ovarian cancer tissues

Given the tissue-specific patterns of enrichment in active regulatory states, we restricted these analyses to regions only marked by H3K27Ac, the most widely profiled marks in Roadmap Epigenomics and ENCODE tissues. We also included in these analyses data we have generated through H3K27Ac-ChIP-seq profiling of primary tissues or cell lines for 26 ovarian cancers representing the different histotypes of invasive disease, and 6 normal cell lines representing putative cells of origin of the different ovarian cancer histotypes (see Methods) (Supplementary Table 3).

We observed enrichment of EOC risk SNPs in H3K27Ac peaks in 38 of the 98 cell types from in Roadmap Epigenomics/ENCODE, and depletion in only 10 cell types (Track 1 of Figure 3 and Supplementary Table 6). EOC risk SNPs were most enriched in H3K27Ac in blood and T-cell tissues and were significantly depleted in all seven brain cell types.

**Figure 3.**
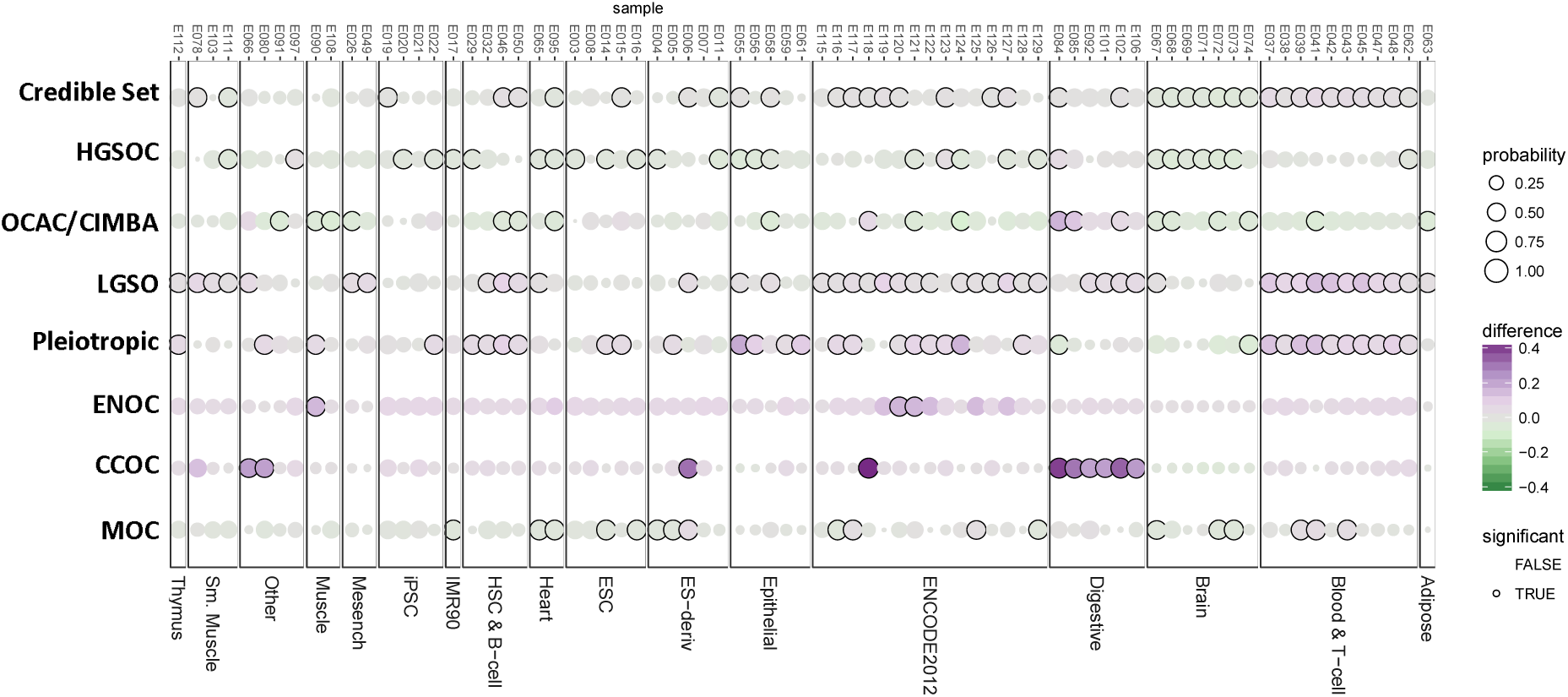
Histotype specific credible causal variants show different patterns of enrichment. Enrichment analyses were performed for each EOC histotype in active regulatory regions marked by H3K27Ac in Roadmap Epigenomics and ENCODE tissues and cell lines. Enriched tissues are shown in purple, depleted tissues in green, and non-significantly enriched tissues in grey. The size of the circle indicates the degree of confidence.

After stratifying EOC risk SNPs by histological subtype, we found the strongest enrichment for risk variants at the 17q12 risk locus for the CCOC histotype; all 8 candidate causal SNPs at this locus lie in intronic regions of *HNF1B* gene (hepatocyte nuclear factor 1 homeobox B) (Figure 3 and Supplementary Table 6), with the greatest enrichment in digestive (E106, E102, E101, E092, E085, E084) and liver (E080) tissues.

We next performed the same analysis for H3K27Ac marks profiled in 38 ovarian cancer related tissues, including ovarian tumors for different histotypes, normal ovarian cancer precursor cell types and data from profiling of whole ovary specimens ^55^. We also compared these data to enrichment for other tissue types from Roadmap Epigenomics/ENCODE which may indicate other tissues of origin for ovarian cancers (e.g. mucinous ovarian cancers, which may arise from cells of the digestive tract). We observed enrichment of EOC risk SNPs across all ovarian tissues except for whole ovary. The strongest enrichment was observed in H3K27Ac peaks in primary HGSOCs in which 197/1432 SNPs (13.75%) overlapped H3K27Ac peaks, compared to 5.6% of the background (control SNPs) (probability > 0.999) (Figure 4a, Supplemental Table 8, and Supplemental Table 9). In parallel, we also estimated enrichment of heritability in these H3K27Ac marks based on common SNPs with similar findings (Supplementary Materials, ‘Enrichment of common SNPs in ovarian cancer related H3K27Ac peaks based on partitioned heritability’ paragraph).

**Figure 4.**
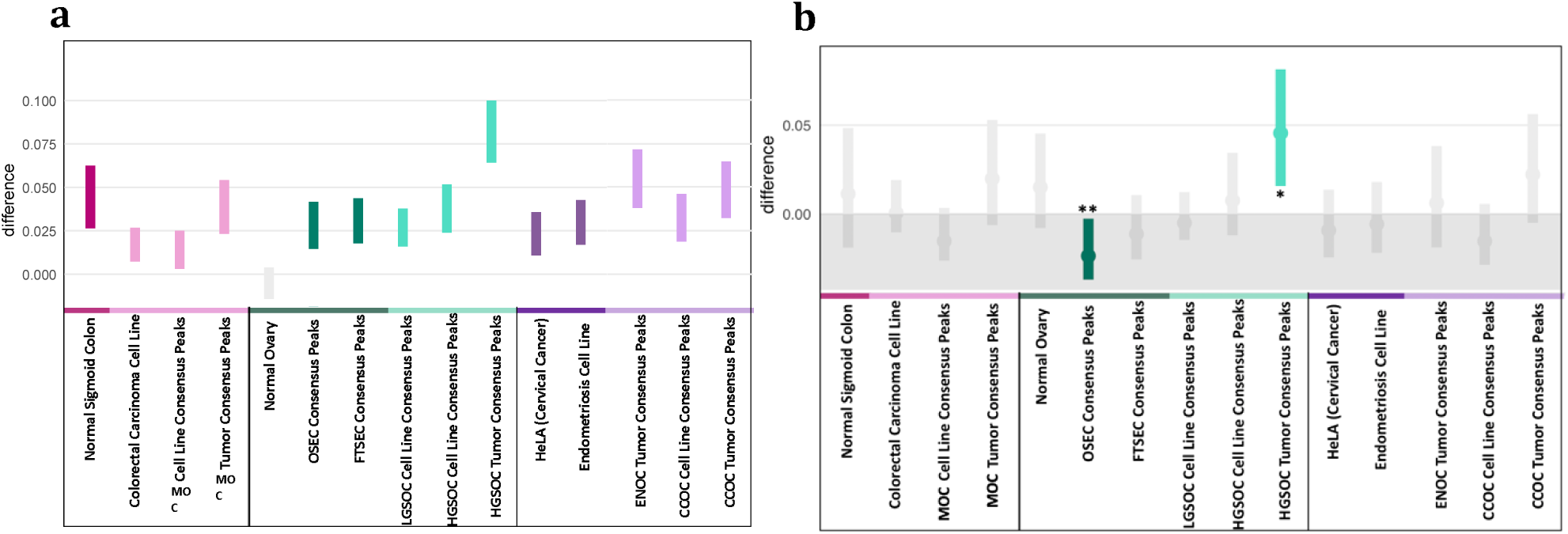
Enrichment of EOC risk variants in ovarian cancer associated tissues and cell lines. (a) EOC credible causal SNPs are significantly enriched in precursor (dark colors) and cell line models of EOC, and primary EOC tumors (light colors). (b) Credible causal SNPs associated with HGSOC are enriched in active regulatory regions in primary HGSOCs (*) and significantly depleted in ovarian surface epithelial cells (OSEC consensus peaks) (**)

We repeated these analyses after stratifying the panel of candidate causal EOC risk SNPs by histotype. In total there were 315 candidate causal risk SNPs specific to HGSOC, 353 SNPs specific to LGSOC, 8 SNPs specific to CCOC, 8 SNPs specific to EnOC, 296 SNPs specific to MOC and 47 SNPs specific to LMP histotypes. Risk SNPs for HGSOC were most significantly enriched in H3K27Ac marks in primary HGSOC tumors; 31/315 (9.8%) risk variants for HGSOC intersect H3K27Ac marks in primary HGSOCs, compared to local background SNPs (difference=0.045, probability=0.999; Figure 4b). Notably, we observed little or no enrichment for HGSOC risk SNPs in H3K27Ac marks generated in HGSOC cell lines, nor in normal FTSECs which are the reported precursors of HGSOC (Figure 4b). HGSOC risk SNPs were also significantly depleted in normal ovarian surface epithelial cells (OSECs). We also observed significant enrichment of risk variants associated with the LMP histotype in H3K27Ac marks in OSECs (Supplementary Tables 10 and 11; Supplementary Figure 2), but no tissue specific enrichments for risk SNPs for other histotypes, which could largely be due to the lack of statistical power to detect enrichment.

### In silico analysis of EOC risk SNPs intersecting transcription factor binding site (TFBS) motifs

We evaluated the putative effects of the 590 EOC risk SNPs intersecting H3K27Ac marks on binding to TFBS motifs using statistical tool, motifbreakR ^63^. The 590 EOC risk SNPs were selected by intersecting with at least one H3K27Ac peak in any of the precursor normal or ovarian cancer cell lines or tumors. 469 out of 590 SNPs were predicted to significantly disrupt at least one TFBS (P-value < 1×10^−5^; Supplementary Table 12), compared to background SNP set which was drawn from credible causal SNPs that did not intersect any EOC-related H3K27Ac marks. Eighty-two SNPs were predicted to break a single TFBS; the remaining SNPs break two or more (on average four) motifs with 5 SNPs predicted to break more than 20 motifs (Figure 5a). At the 18q11.2 locus, which confers risk of HGSOC, rs9955681 located in an intron of the *LAMA3* gene, was predicted to break 67 different motifs; and at the 4q26 EOC locus, rs7671665, which is located in intron 2 of the *SYNPO2* gene was predicted to break 31 different motifs (Supplementary Table 12, Figure 5a).

**Figure 5.**
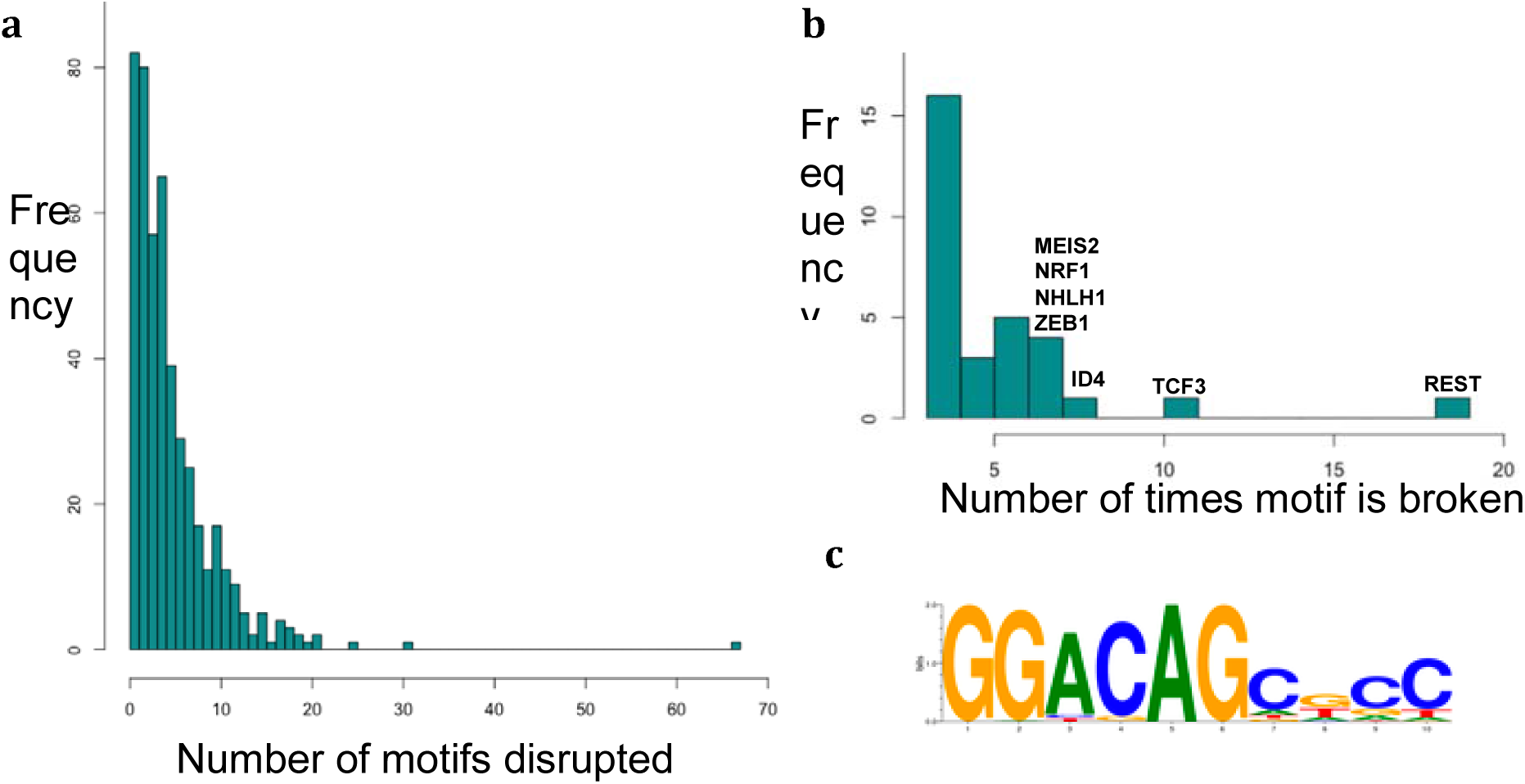
EOC risk SNPs disrupt TF motifs at risk loci. (a) Number of motifs disrupted by credible causal SNPs intersecting EOC-related H3K27Ac peaks. (b) Number of times motif is broken by credible causal SNPs that overlap EOC-related H3K27Ac peaks. (c) REST motif logo from motifbreakR.

The most frequently disrupted TFBS motifs were for REST (repressor element-1 silencing transcription factor) disrupted by 19 SNPs across 12 loci (P-Value = 0.0028); TCF3 (Transcription factor 3) disrupted by 11 SNPs (P-Value = 0.0075); and ID4 (DNA-binding protein inhibitor), which was disrupted by 8 SNPs (P-Value = 0.0025) (Figure 5b and Supplementary Table 13). The motif for the epithelial-specific transcription factor EHF, which is overexpressed in EOC tumors, induces apoptosis and impairs cell adhesion and invasion after knockdown in EOC cell lines ^73^ was broken by 6 SNPs at five EOC risk loci associated serous and mucinous histotypes (1p36, 2q13, 2q31, 8q24, 19p13).

## Discussion

Identifying the functional effects of common susceptibility variants identified by GWAS on is an important step in delineating the biological mechanisms underlying disease and in understanding the earliest stages of disease pathogenesis. In this study, we examined the heritability for risk variants associated across all ovarian cancer and for each of each of the different histotypes of disease. Moreover, we partitioned heritability into broad functional categories to identify those that that are the drivers of neoplastic initiation and progression.

We identified enrichment of EOC credible causal SNPs into active regulatory elements marked by H3K27Ac in ENCODE and Roadmap Epigenomics public datasets. This indicated germline risk variants that contribute to disease biology via disruption of enhancer activity in cell and tissue specific active regulatory regions, rather than regulatory elements that are active across a broad range of cell types. We further identified strong enrichment of the full credible causal variant list in 14 of the 15 highly EOC relevant cell types included. We observed clear patterns of enrichment of HGSOC germline risk SNPs in HGSOC tumors, and depletion of these variants in H3K27Ac from precursor normal cells. These findings suggest that HGSOC germline risk variants affect cancer progression or development rather than initiation, and underscore the need for variant annotation using cell types relevant to disease. Finally, we identified TFs whose binding motifs are significantly disrupted by EOC risk SNPs in active regions.

The cells of origin for the different histotypes of ovarian cancers are not precisely known. Fallopian tube epithelial cells are the most like precursors of HGSOCs and CCOC and EnOC are more likely arise from endometriosis ^4–8^. Our comprehensive H3K27Ac ChIP-seq data in ovarian and non-ovarian cancer tissues makes it possible to identify the putative cells of origin of disease. The significant depletion of HGSOC credible causal variants we observed in H3K27Ac from OSECs active regions (Figure 4b) is consistent with an emerging consensus that HGSOC is less likely to arise from ovarian surface epithelial cells ^4,5,74^. The significant enrichment of LMP risk variants in OSECs active regions supports a role for this cell type in this histotype (Supplementary Table 10 and Supplementary Figure 2) ^75,76^.

It has been hypothesized, with supporting data from pathology examination ^76^, that ovarian surface epithelium invaginates into the underlying stroma of the ovary to form inclusion cysts that undergo transformation to become malignant ^76^. LMP and LGSOC are likely to arise from transformed OSECs trapped within inclusion cysts ^75^ and the significant enrichment of six SNPs at two LMP rick loci (4q32.2 and 5p15) in OSECs (Supplementary Table 11) supports an OSEC origin for these tumor types.

CCOCs are strongly associated with endometriosis, and may derive from ciliated epithelial cells in ovarian endometriosis lesions ^77,78^. Only one locus has been confirmed to be associated with CCOC risk (the *HNF1B* 17q12 locus) which makes it challenging to investigate the likely cells of origin in the current study. We observed a strong enrichment for CCOC credible causal variants at this locus in digestive and liver cells which supports this (Track 7 of Figure 3). This locus is pleiotropic for both HGSOC and CCOC, but we only observed significant enrichment in H3K27Ac marks for CCOC and MOC tumors and cell lines (Supplementary Figure 4a). Here all 8 candidate causal SNPs at 17q12 lie in intronic regions of the *HNF1B* gene. *HNF1B* has been reported as a susceptibility gene and is highly expressed in CCOCs but largely absent in HGSOCs ^7,79^. We further investigated gene expression of *HNF1B* across our previously generated ovarian cancer tumors RNA-seq data^41^. We found *HNF1B* is expressed in MOC, EnOC, and CCOC, but not in HGSOC (Supplementary Figure 4b), which is consistent with the difference in H3K27Ac enrichment between histotypes.

We present here an approach to annotate risk SNPs that may influence transcriptional regulation by interacting with the epigenomic landscape to disrupt TF binding and alter gene regulation and expression. For example, SNPs rs7671665 and rs9955681 were predicted to break the greatest number of motifs. We identified SNP rs7671665 that breaks 31 motifs within a regulatory element present in a wide range of Roadmap Epigenomics and ENCODE cell types and most of our panel of EOC related cell types. This SNP is an eQTL located within intron 2 of *SYNPO2*, and is reported to loop to the promoter of *SYNPO2* ^80^ and *METTL14* ^81^, a component of N6-methyladenosine (m6A) methyltransferase complex. This complex controls post translational modification of m6A RNA and has been implicated in cancer, cell differentiation and proliferation in development pathways ^82^. Interestingly, m6A is reported to be enriched in the 3’UTR ^83^, which was the most significantly enriched biofeature in our partitioning of heritability analysis. Another example is SNP rs9955681, which is predicted to break 67 TF motifs in EOC tumors active regions. This SNP is located in an intron of *LAMA3*, a known enhancer in breast and cervical cancer cell lines and gastrointestinal tissues ^29,33^. This SNP is also a known eQTL in previous HGSOC susceptibility gene analyses ^84^.

In conclusion, we have applied enrichment approaches to identify overrepresentations of risk SNPs within specific biofeatures. By intersecting risk SNPs with a catalogue of regulatory elements, we identify putative enhancers impacted by risk variants that help explain the underlying functional mechanisms mediating genetic risk as ovarian cancer susceptibility loci. In additional we have shown the power of these approaches to elucidate the putative cells of origin of the different ovarian cancer histotypes, providing support for previously known cell types, and identifying other novel cell types associated with other histotypes. Finally, these studies have defined sets of putative causal variants at ovarian cancer risk loci, that warant further functional analysis to identify the genetic and regulatory mechanisms that drive initiation and early stage development of ovarian cancers.

