## Supplemental Methods for "Ovarian Cancer Risk Variants are Enriched in Histotype-Specific Enhancers that Disrupt Transcription Factor Binding Sites"

### Supplementary Materials

**Enrichment of common SNPs in ovarian cancer related H3K27Ac peaks based on partitioned heritability**

We have demonstrated that risk SNPs are enriched in the epigenome in a tissue-specific manner, highlighting the need for ovarian cancer associated tissues to improved out understanding of their functional roles in the relevant cell type(s).

We estimated the functional enrichment of the narrow sense heritability for ovarian cancer based on common SNPs, and further demonstrate enrichment of credible causal risk SNPs in active regulatory elements. In parallel to FunciVar functional enrichment test in EOC related tissues, we also estimated functional enrichment of heritability based on common SNPs. HGSOC tumors presented the strongest enrichment in both the partitioning of heritability and enrichment of credible causal risk variants, across all precursor normal and cancer associated tissues and cell lines (15.6-fold enrichment, p=0.005; Supplementary Figure 3a). These analyses suggest that risk variants for EOC are functional in enhancers that are active in tumors, rather than precursor cell types, and may contribute to disease progression more than initiation. This enrichment results could potentially be driven by the fact that HGSOC cases are a large proportion of the cohorts used in EOC GWAS, as it is the most prevalent histotype of EOC, and therefore contribute more to the association signal in analysis that combined patients of different histotypes in a single phenotype.

By stratifying the credible causal SNPs to investigate the enrichment of HGSOC specific risk variants we partitioned the heritability for HGSOC across H3K27Ac peaks, and observed the strongest and most significant enrichment in HGSOC tumors (11.5-fold enrichment, p=0.02; Supplementary Figure 3b). Given that relatively few risk alleles are pleiotropic with other types of cancer, the cell type specific enrichment observed in HGSOC tumors is particularly important.

### Supplementary Figures

**Supplementary Figure 1. Credible Causal SNPs for EOC are found in the non-coding genome.** (a) Annotation of credible causal SNPs for EOC with genic features with SNPnexus. (b) FunSeq2 somatic scores are higher in credible causal SNPs for all EOC (lavender) than the set of combined background SNPs (green) used in SNP enrichment analyses.

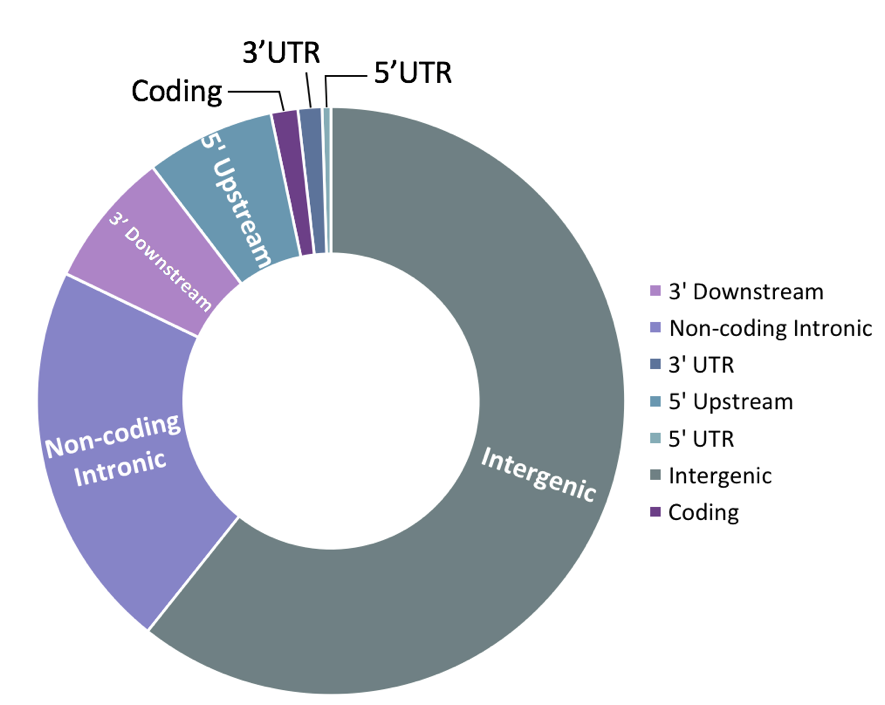

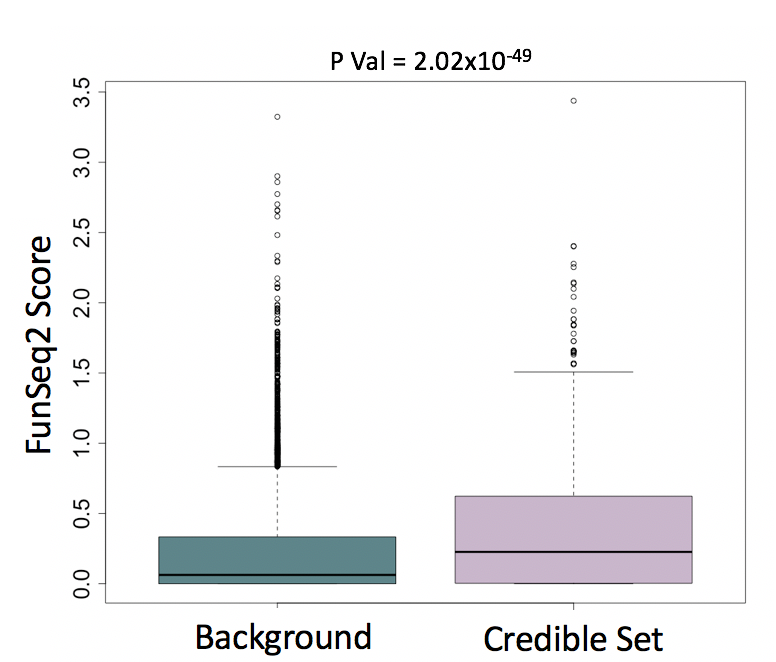

**a**

**b**

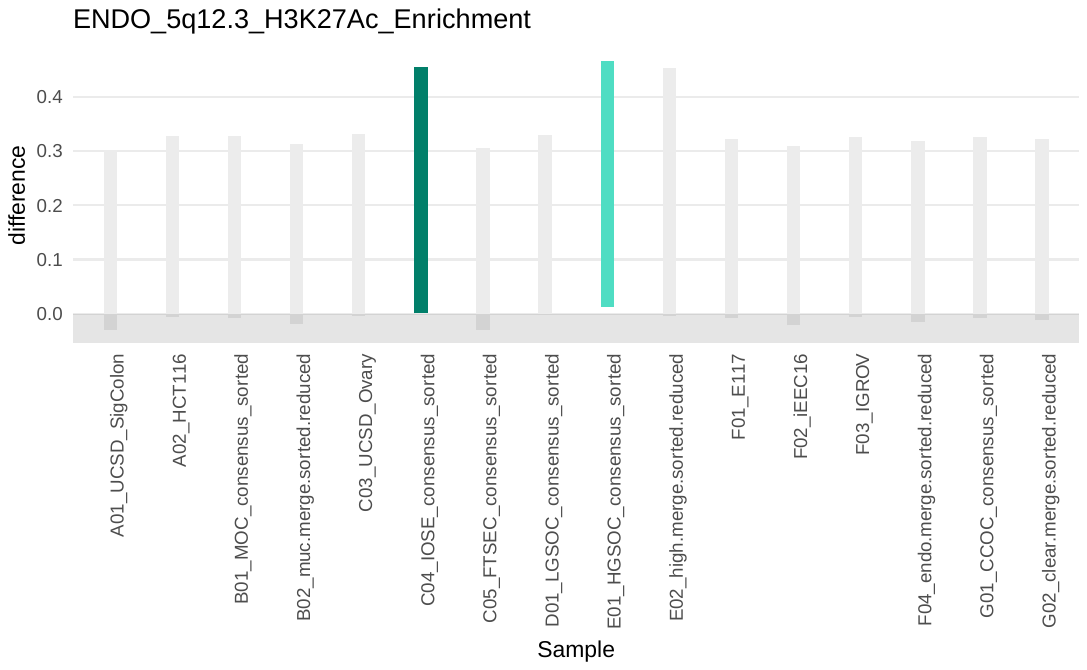

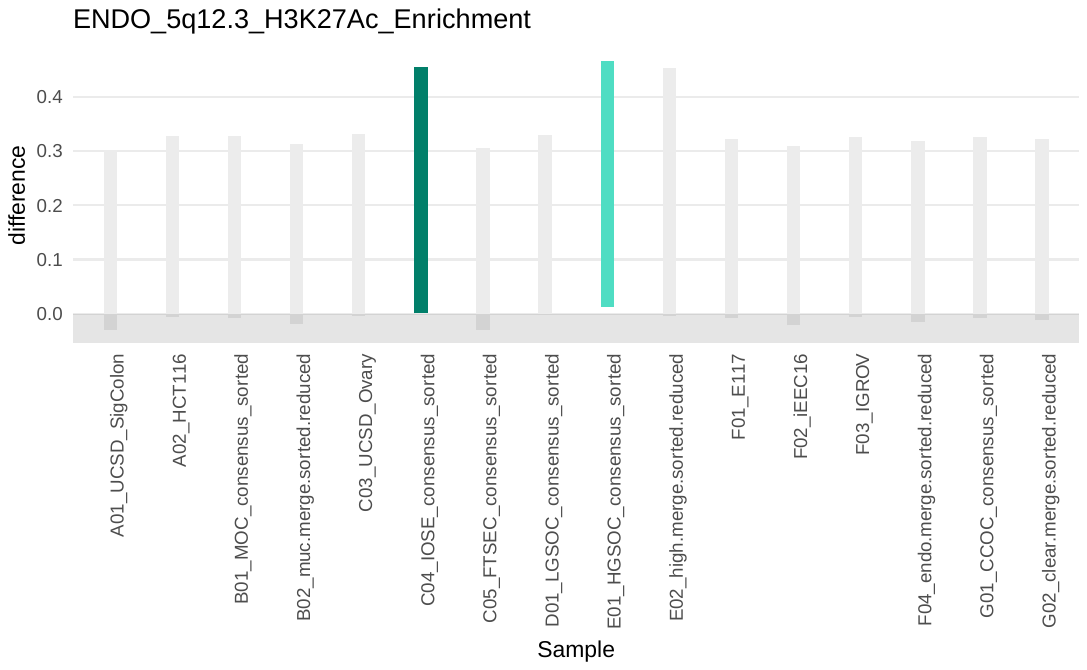

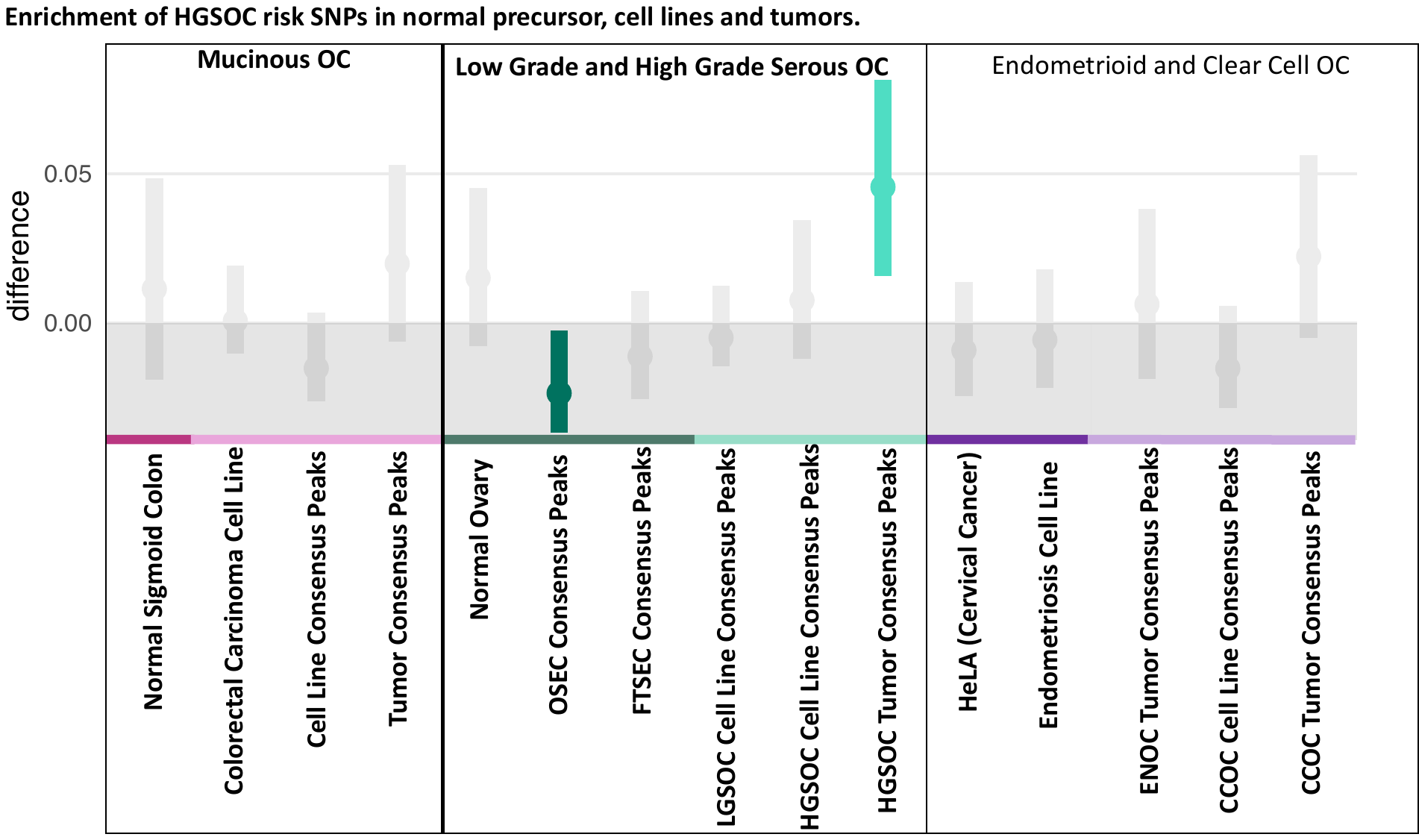

**b) Endometrioid**

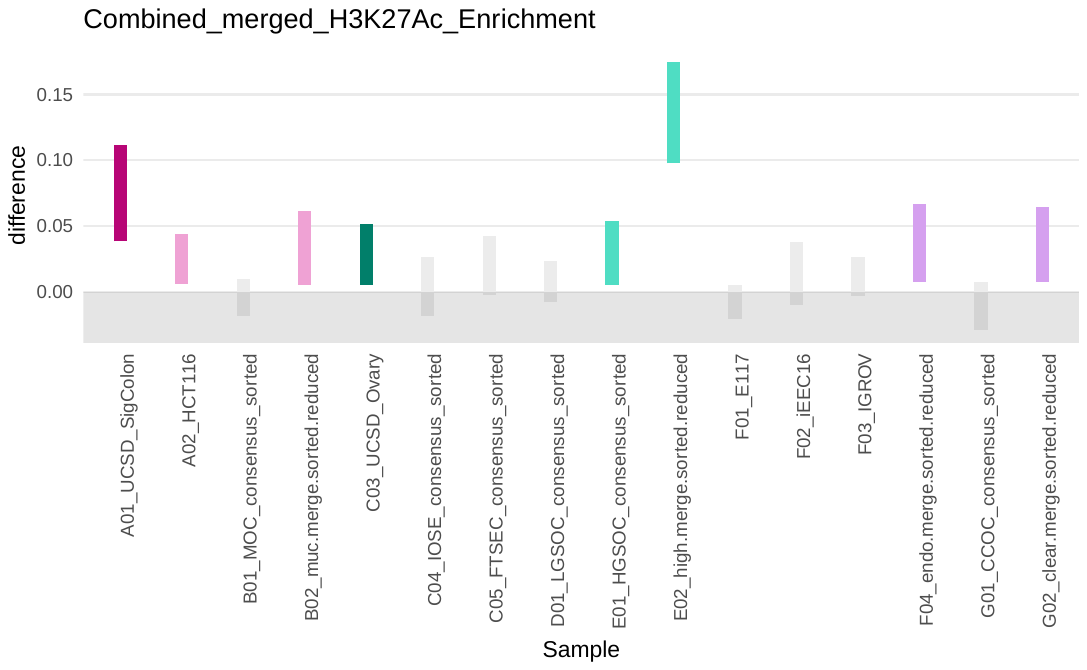

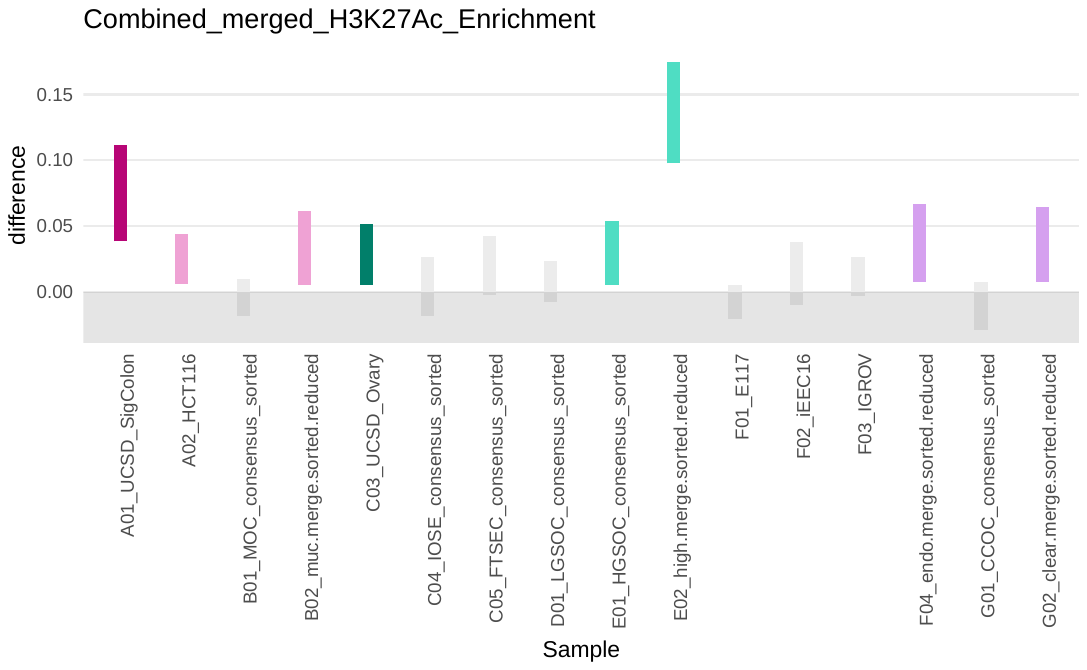

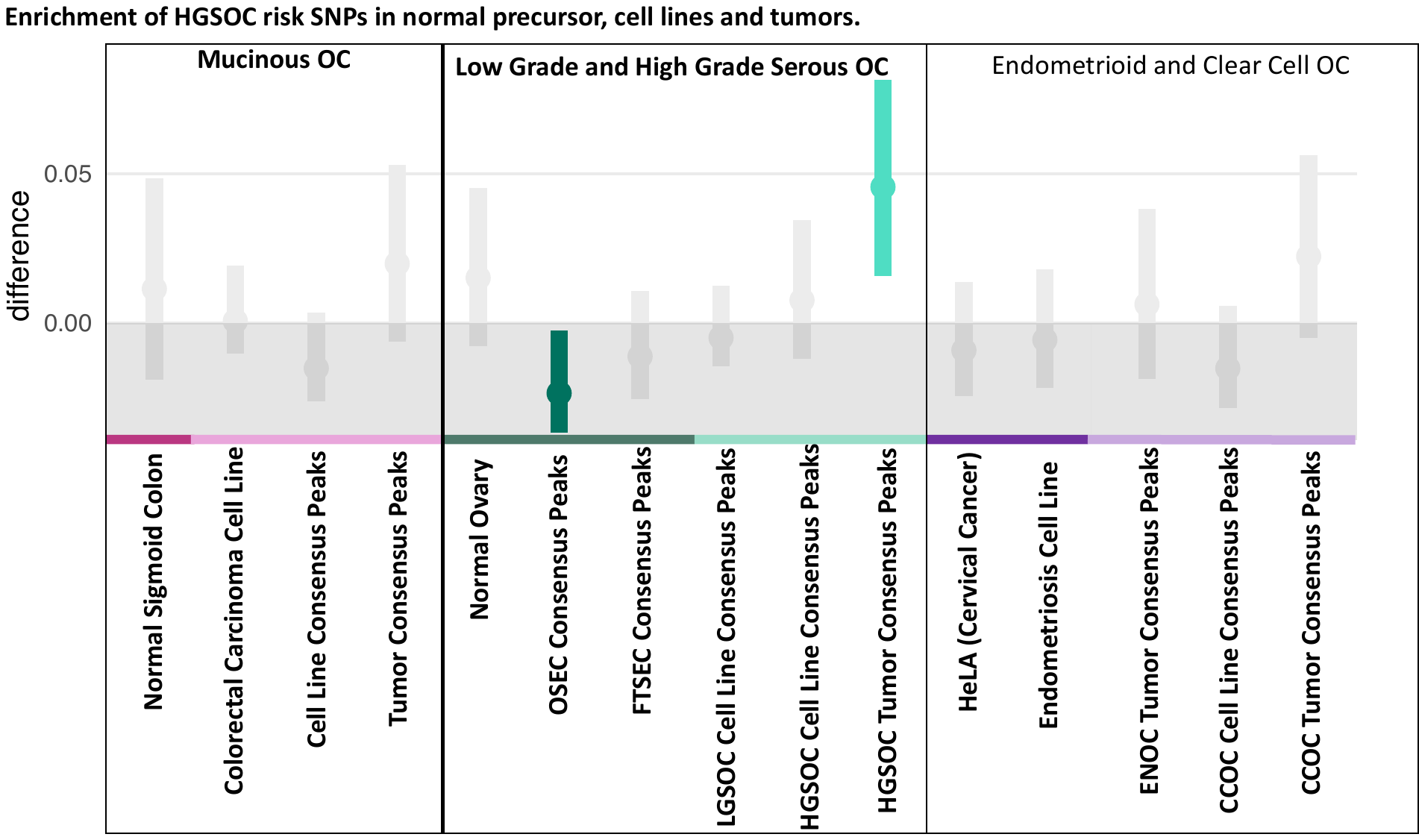

**a) All Combined**

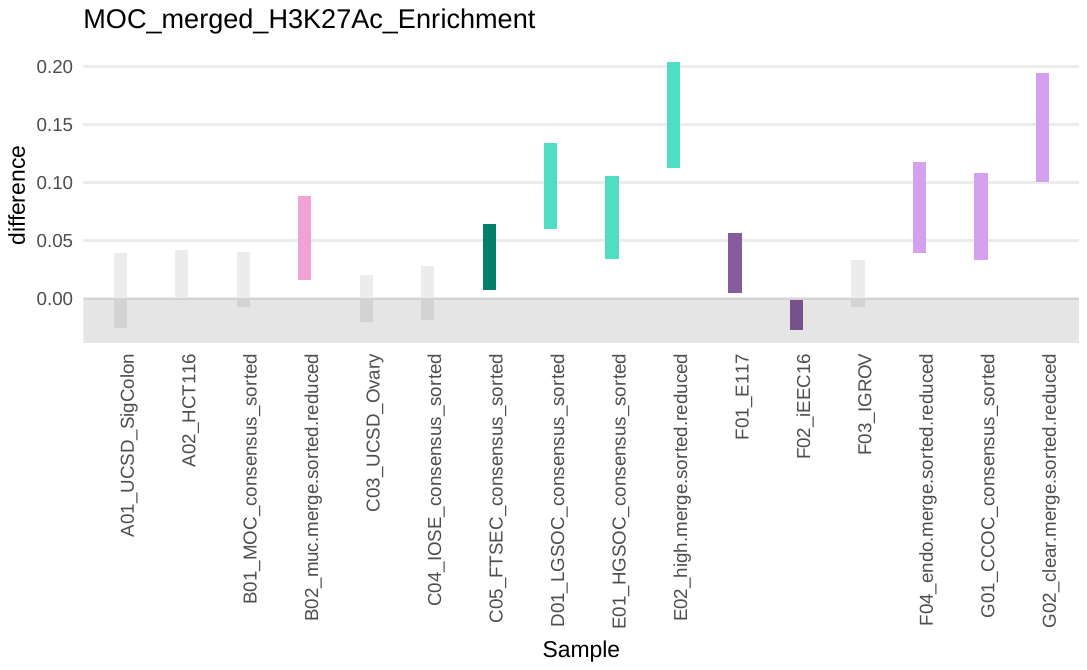

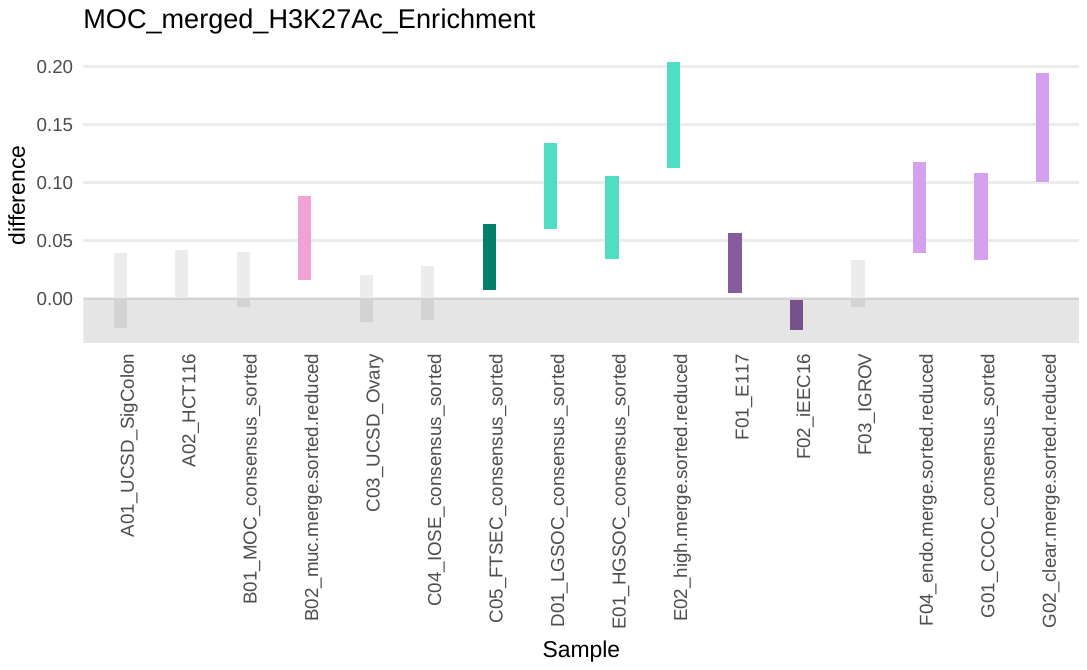

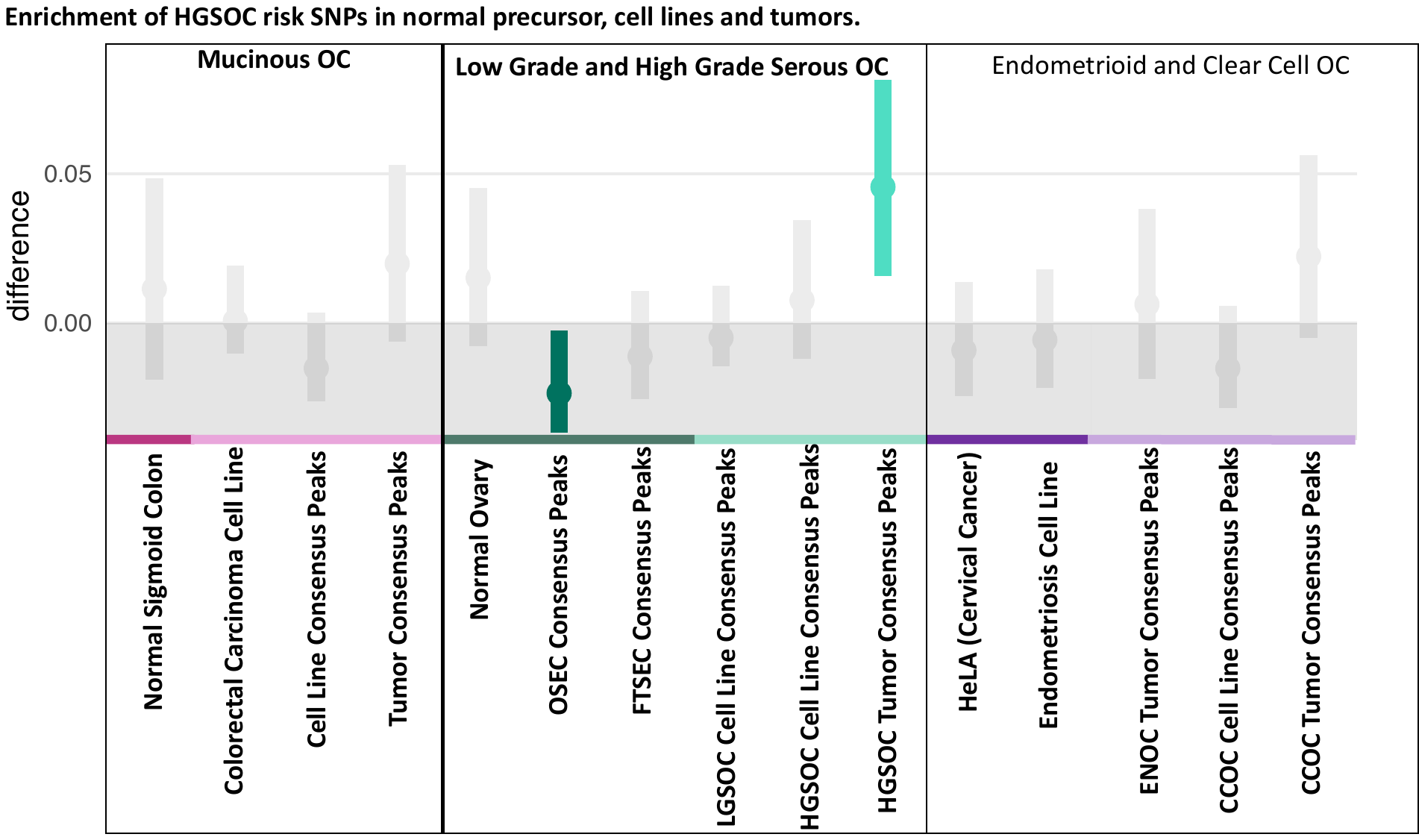

**c) Mucinous**

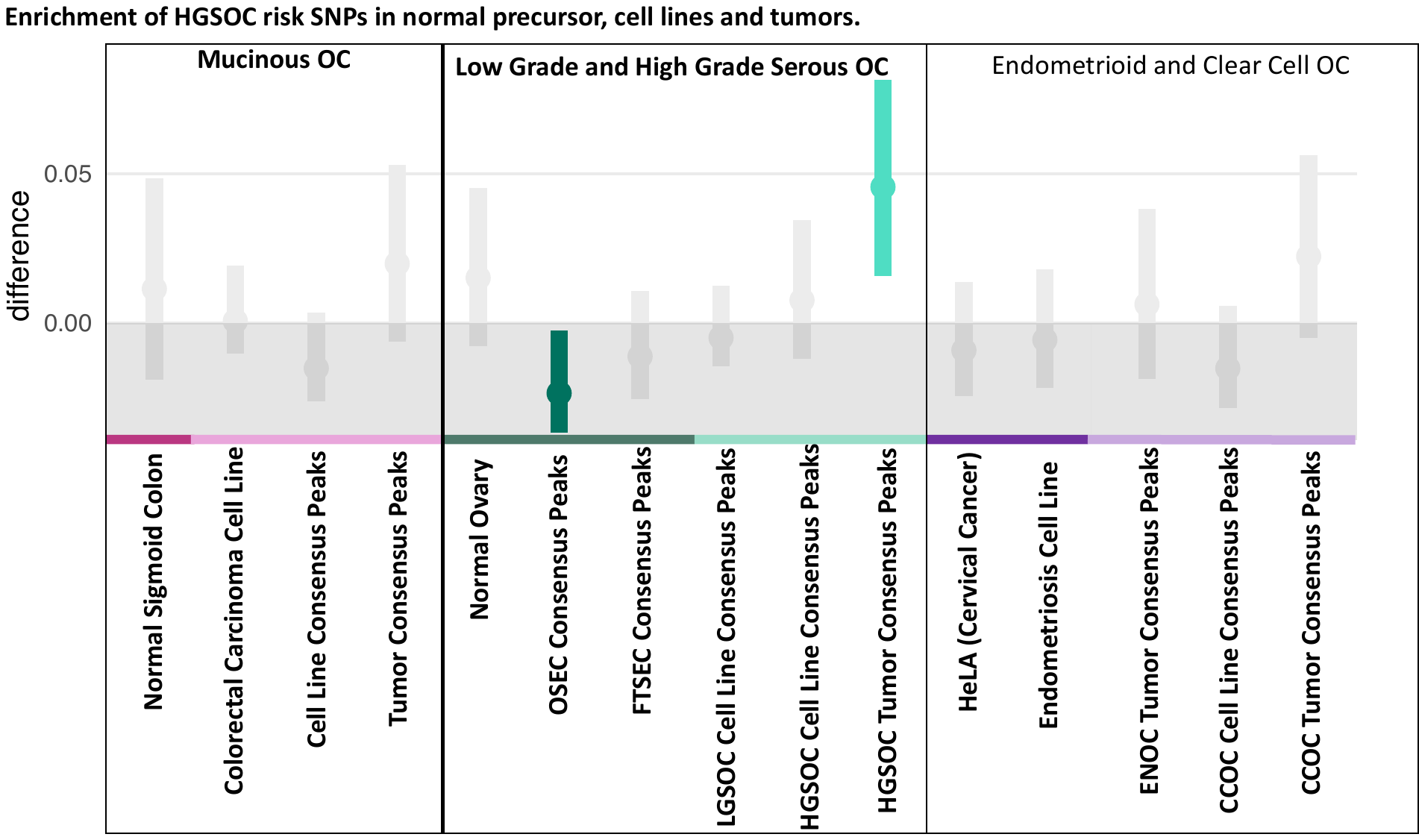

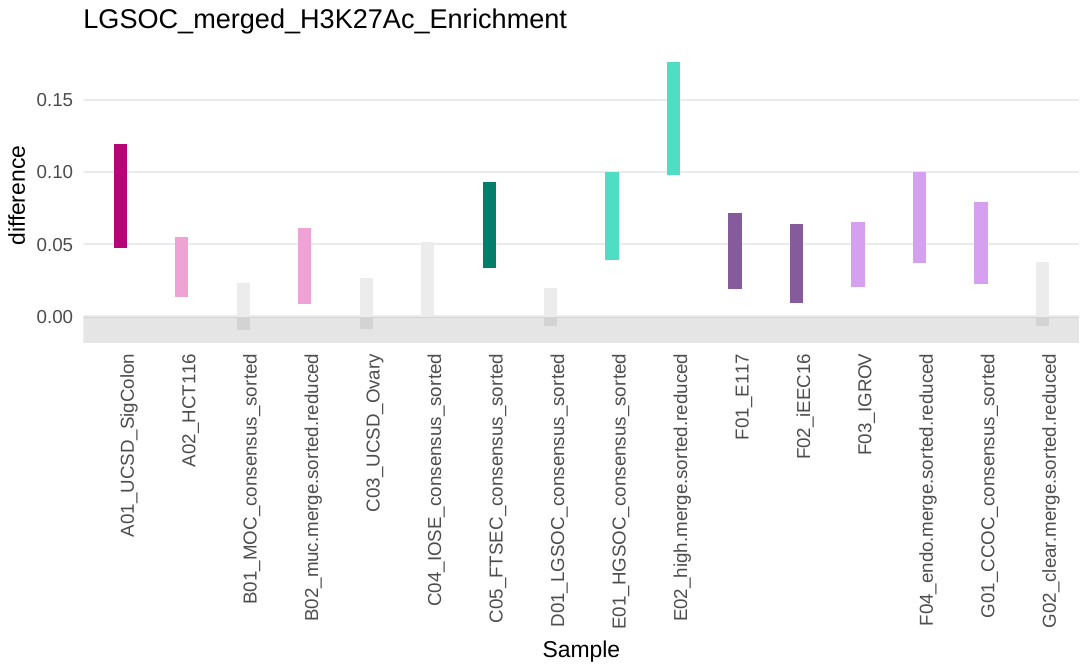

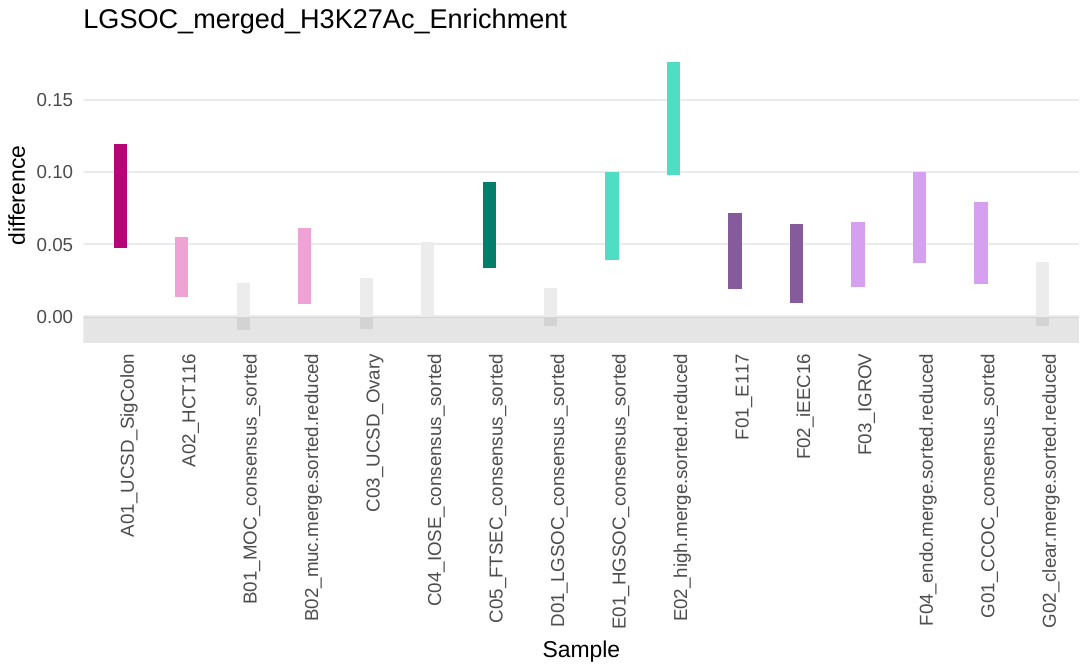

**d) Low Grade Serous**

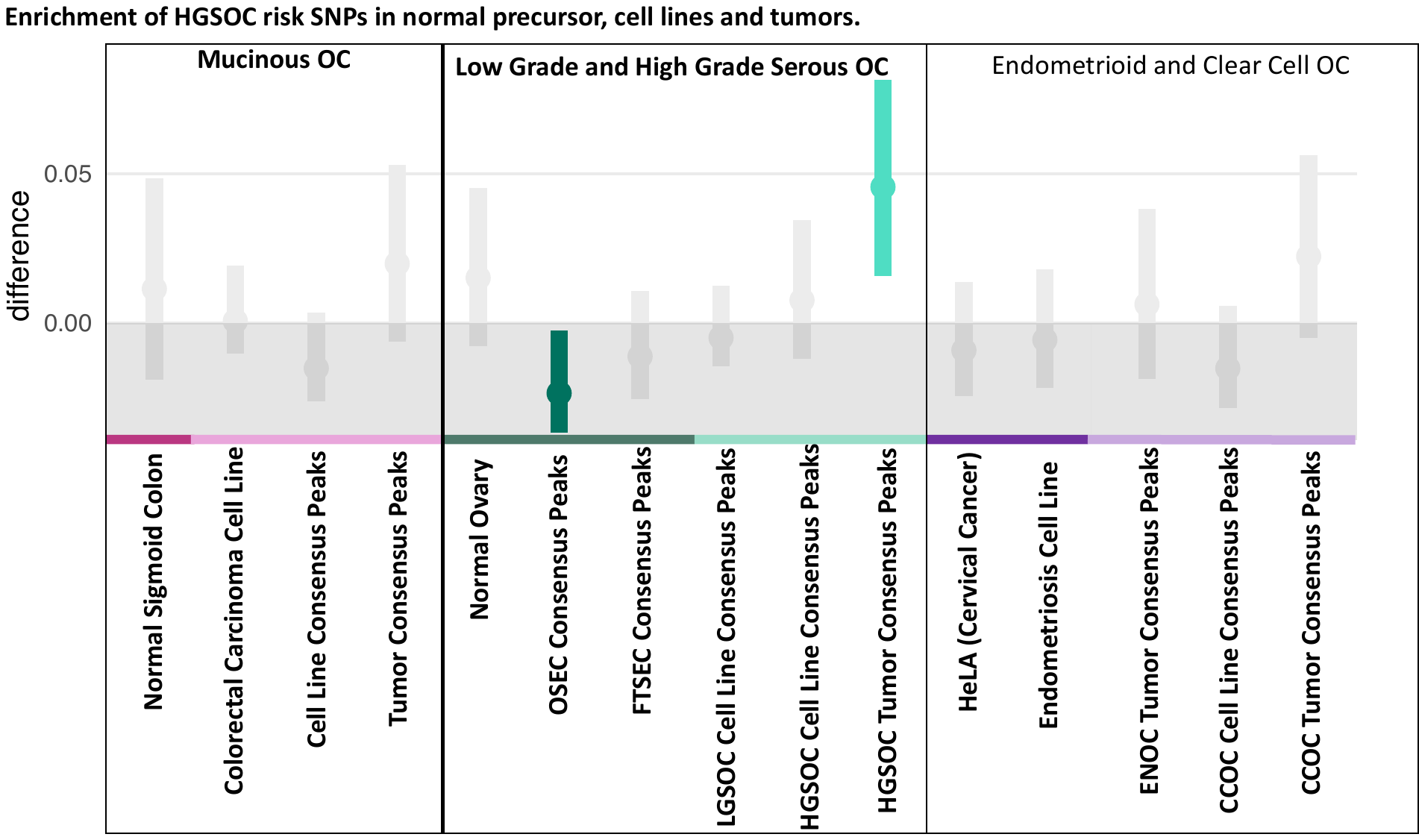

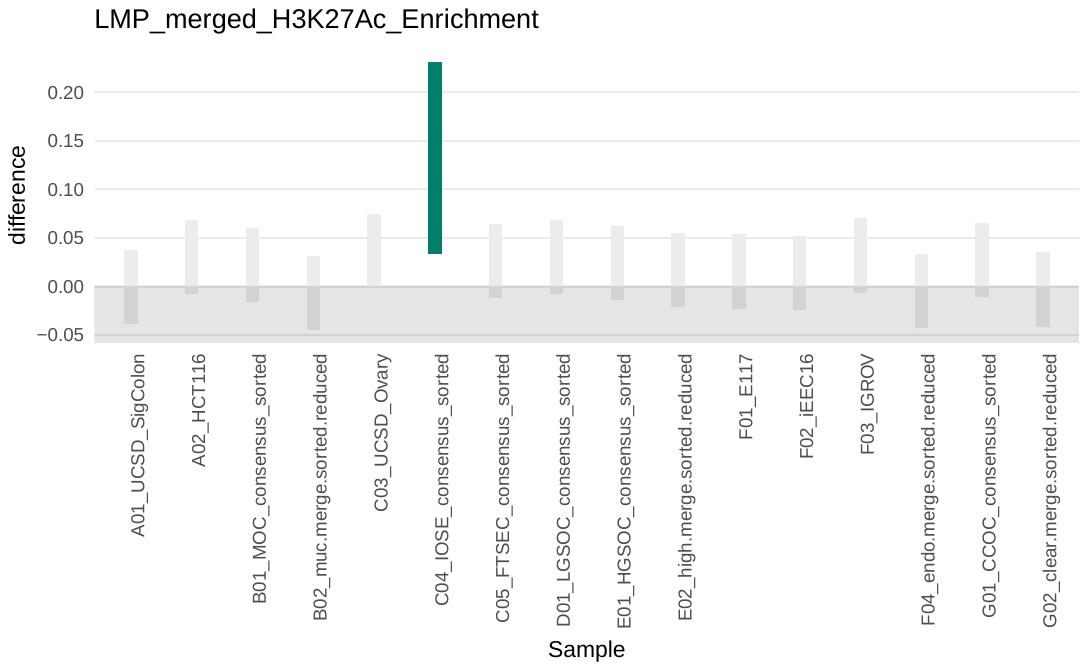

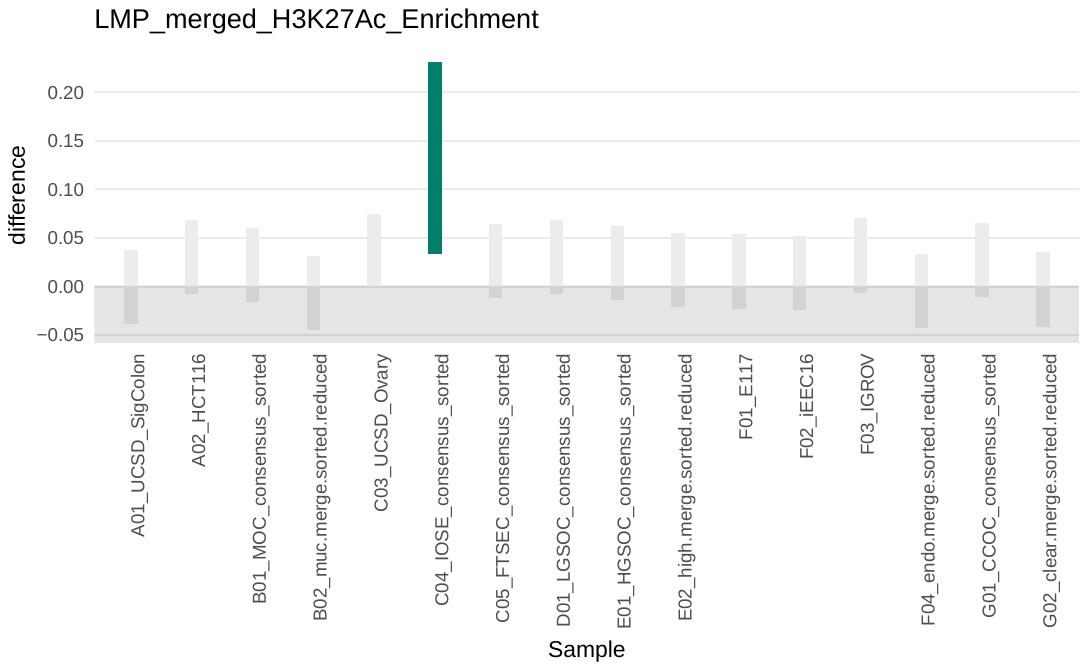

**e) Low Malignant Potential**

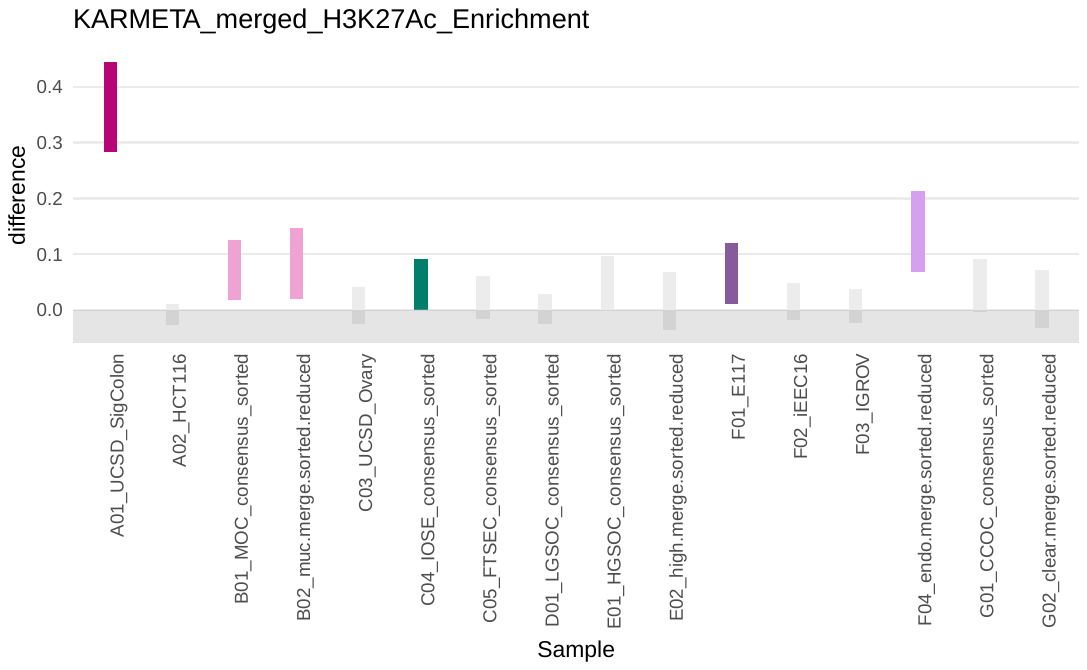

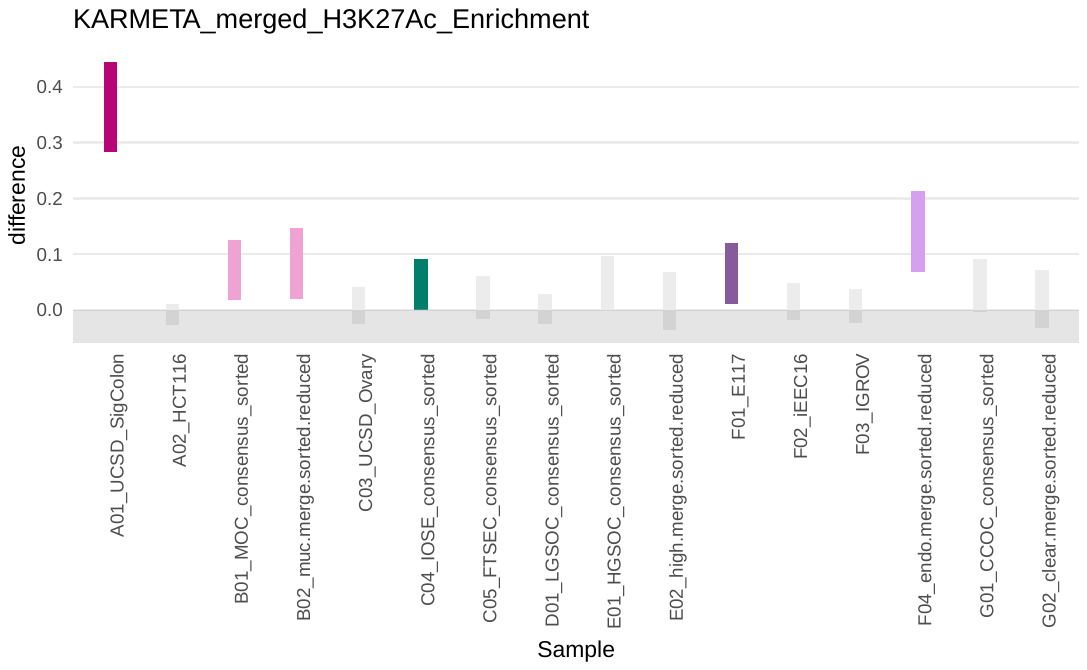

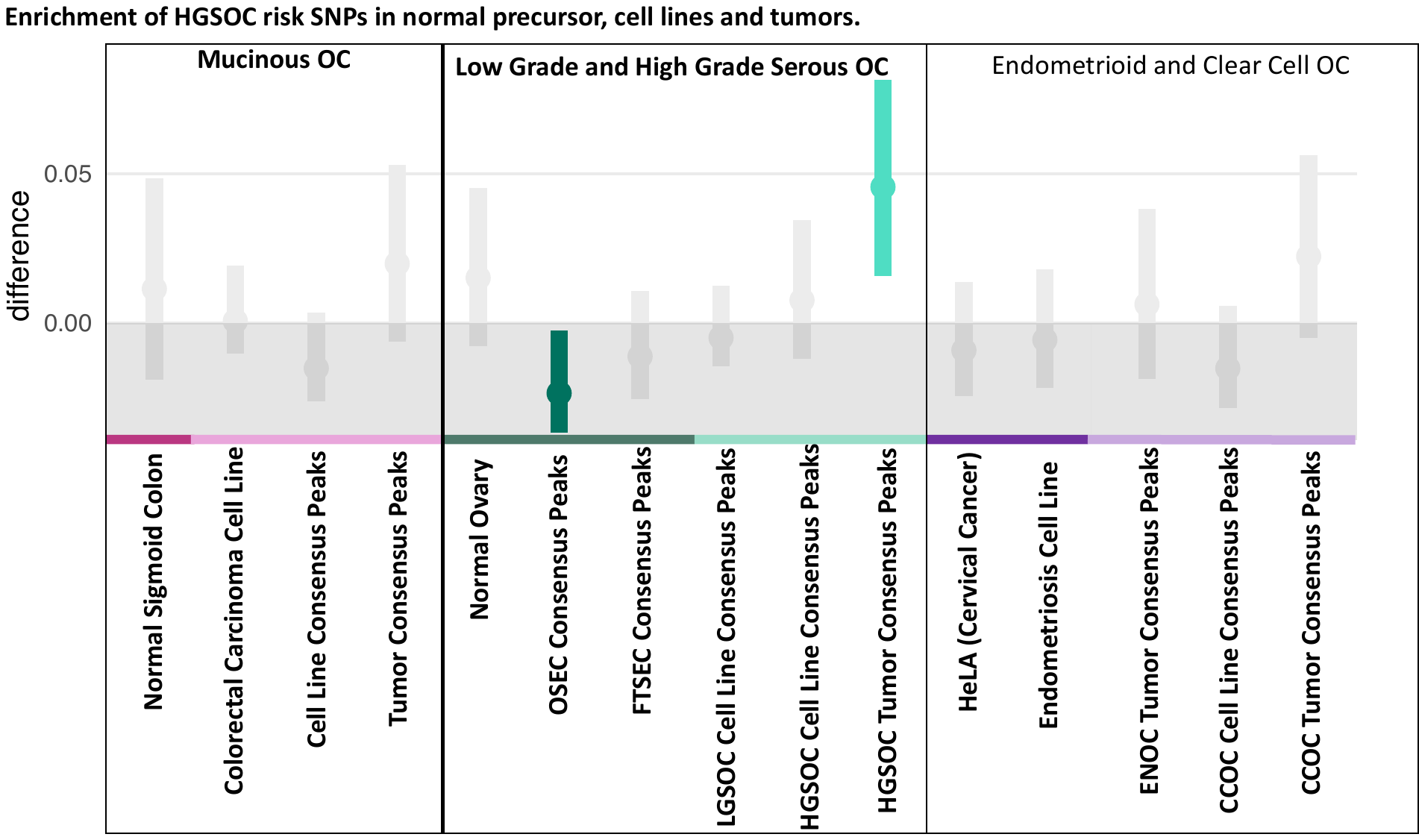

**f) Pleiotropic**

**Supplementary Figure 2. Histotype specific risk variants are enriched in tissues and cell lines related to EOC.**

**Supplementary Figure 3**. Enrichment estimates from GWAS summary statistics were measured as a function of partitioned heritability and proportion of SNPs under the peaks, ${\text{Pr(}h}_{g}^{2})/\text{Pr(}\text{SNPs)}$. Tissues or cell lines with p<0.05 were labelled with star. (a) Enrichment estimates from EOC GWAS summary statistics. (b)Enrichment estimates from HGSOC GWAS summary statistics.

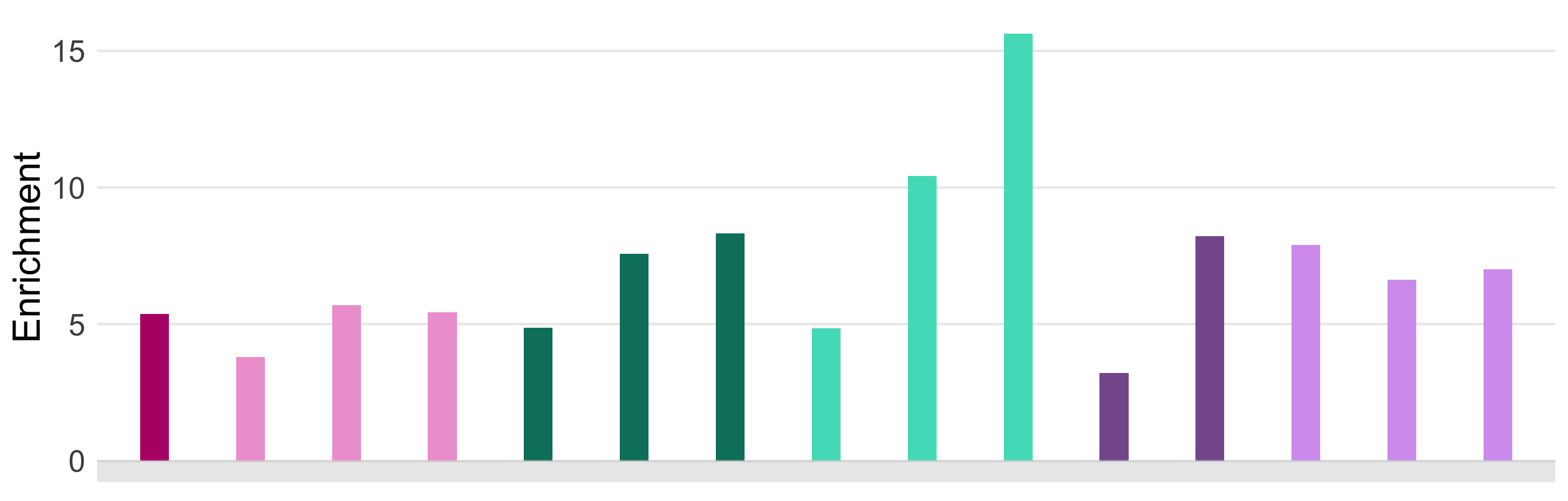

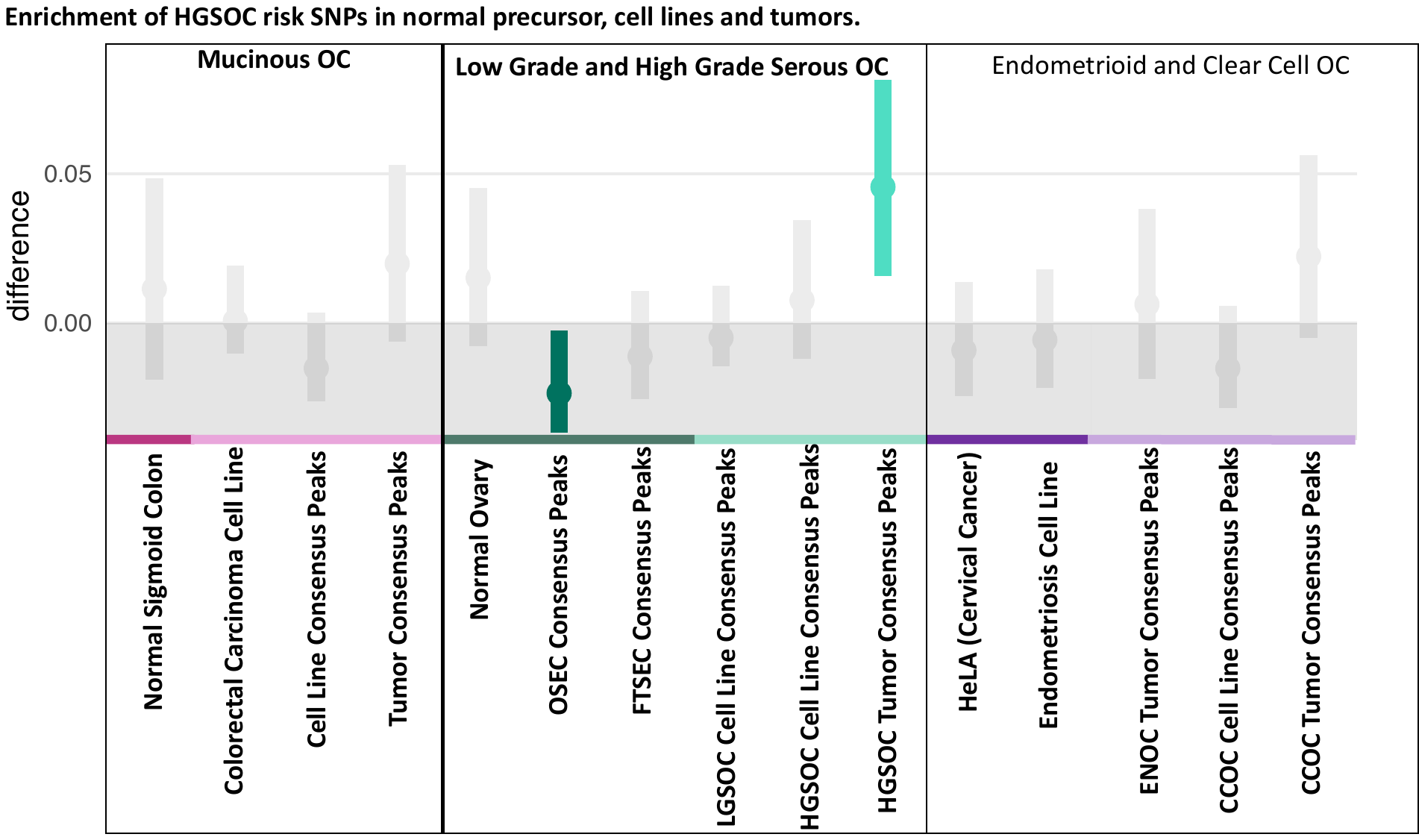

*

*

*

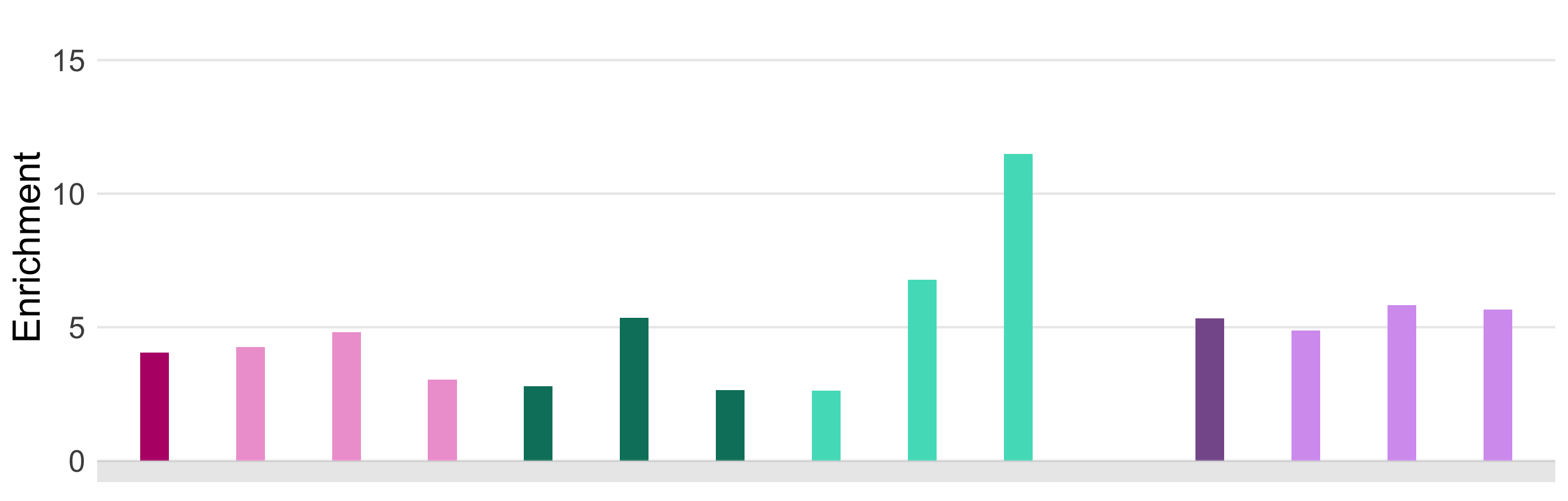

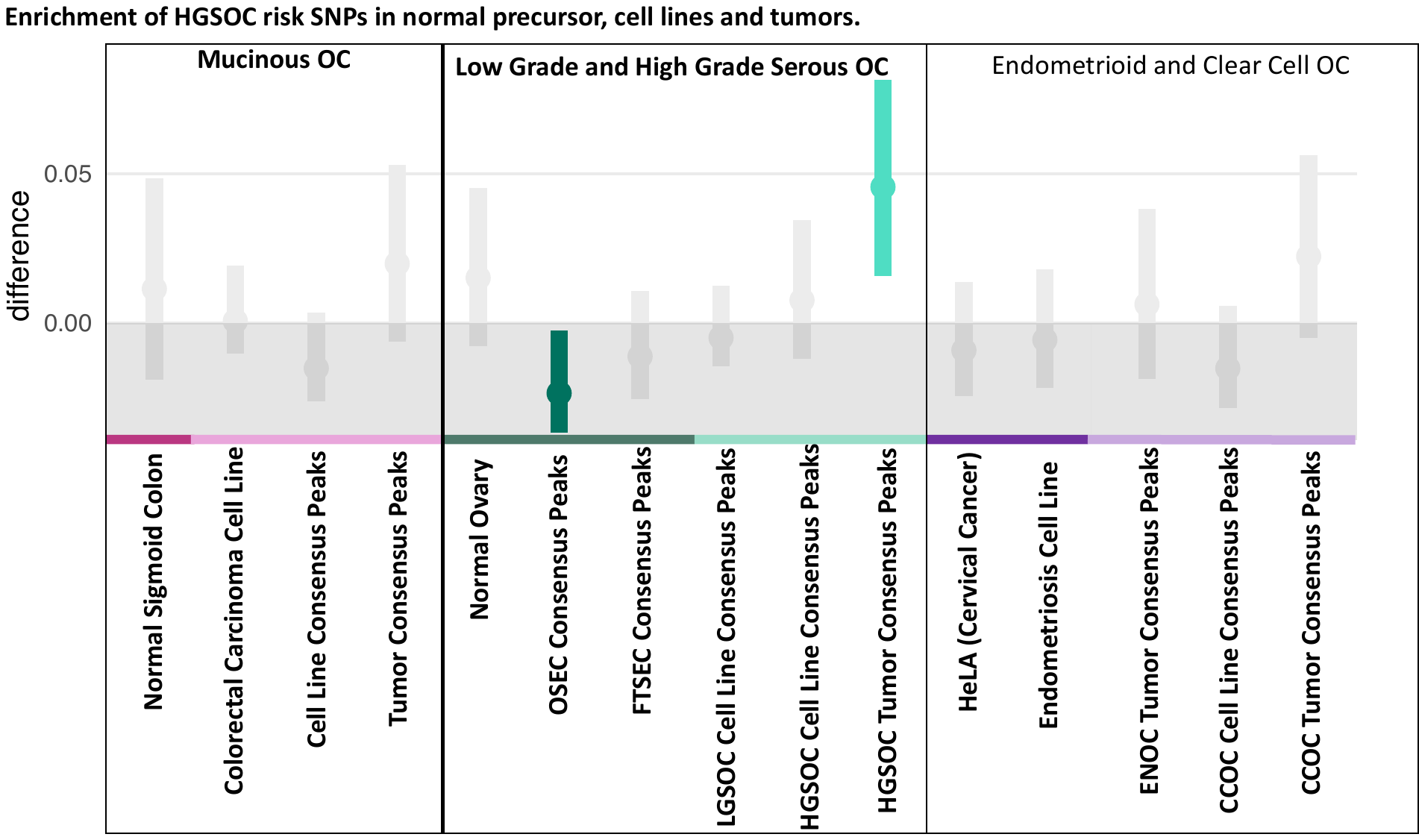

*

**a**

**b**

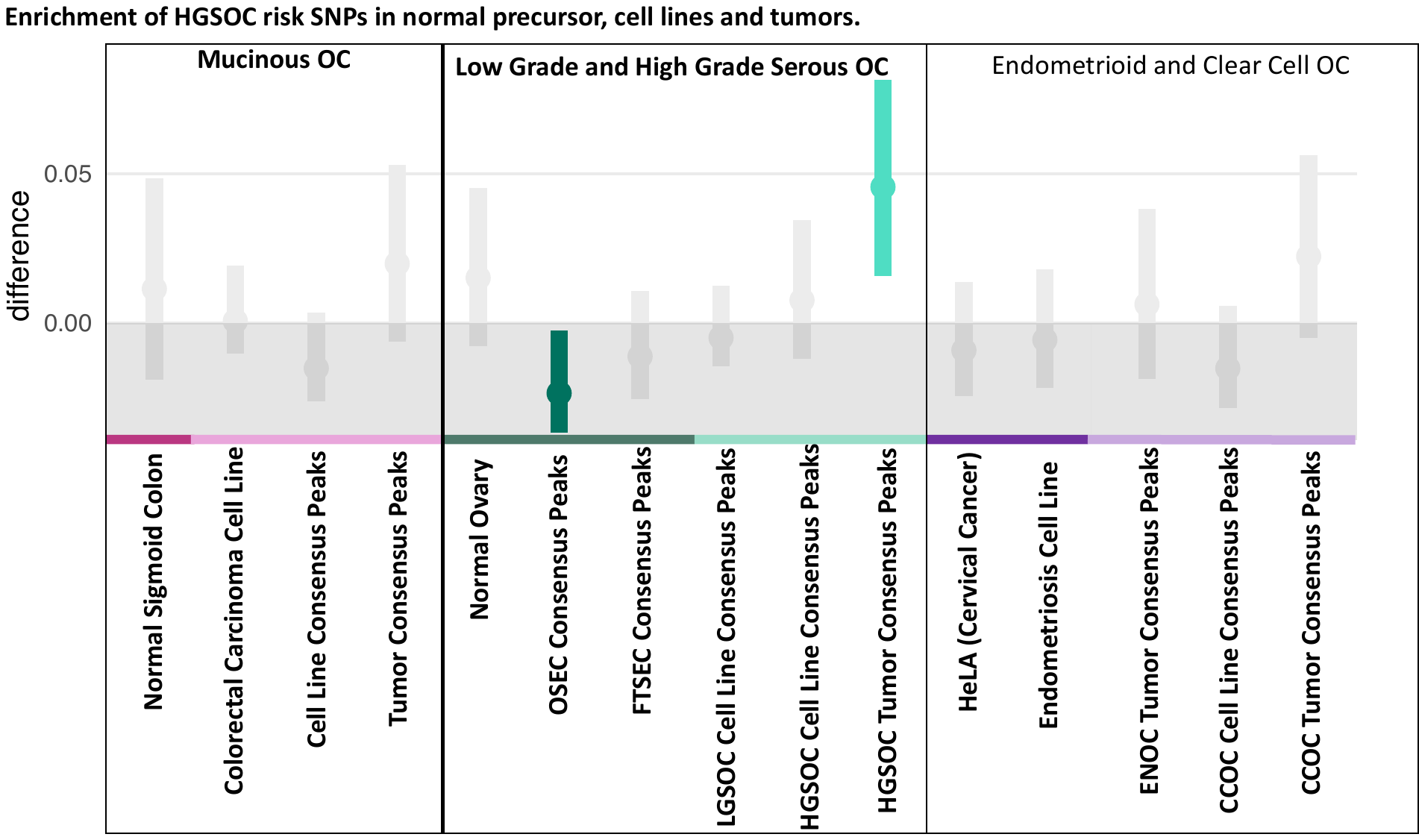

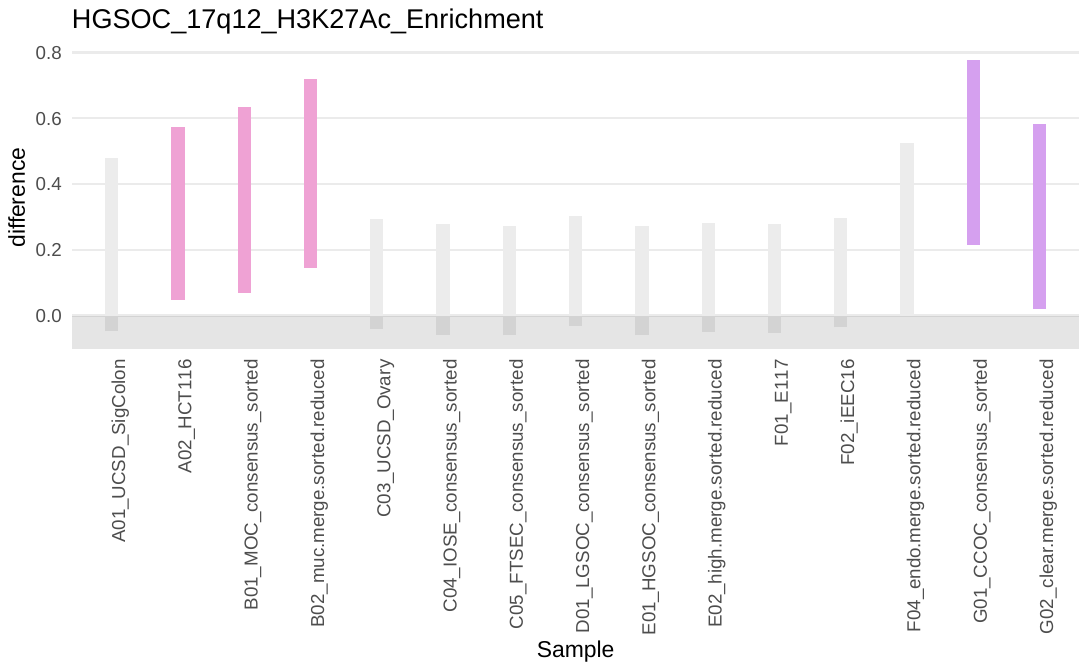

**MOC**

**MOC**

**a**

**b**

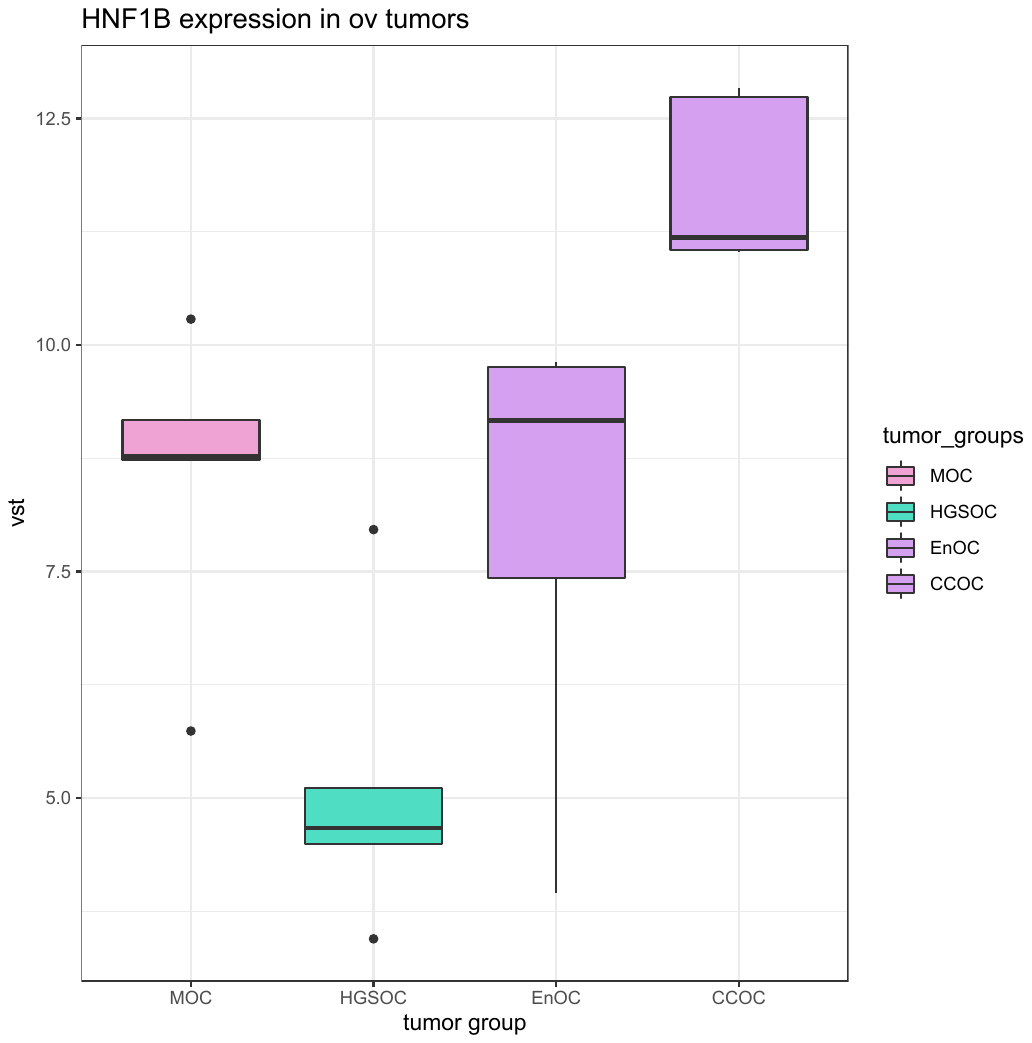

Gene expression level

**Supplementary Figure 4**. (a) CCOC credible causal variants at 17q12 are enrichened in H3K27Ac peaks of MOC and CCOC tumors and cell lines. (b) HNF1B gene is expressed in MOC, EnOC, and CCOC.
